# Rapid redistribution and extensive binding of NANOG and GATA6 at shared regulatory elements underlie specification of divergent cell fates

**DOI:** 10.1101/2021.07.28.454132

**Authors:** Joyce J. Thompson, Daniel J. Lee, Apratim Mitra, Sarah Frail, Ryan Dale, Pedro P. Rocha

## Abstract

Establishment of divergent cell types from a common progenitor requires transcription factors (TFs) to promote lineage-restricted transcriptional programs while suppressing alternative fates. In the mouse blastocyst, cells of the inner cell mass (ICM) coexpress NANOG and GATA6, two TFs that drive the bifurcation of these progenitors into either the epiblast (Epi) or the primitive endoderm (PrE), respectively. Here, using in vitro differentiation, we describe the molecular mechanisms of how GATA6 quickly induces the PrE fate while repressing the Epi lineage. GATA6 functions as a pioneer TF by inducing nucleosome repositioning at regulatory elements controlling PrE genes, making them accessible for deposition of active histone marks and leading to rewiring of chromatin interactions and ultimately transcriptional activation. GATA6 also binds most regulatory elements of Epi genes followed by eviction of the Epi-specific TFs NANOG and SOX2, loss of active histone marks, and reduction in chromatin accessibility that culminates in transcriptional repression. Unexpectedly, evicted NANOG and SOX2 transiently bind PrE regulatory elements occupied by GATA6. Our study shows that GATA6 binds and modulate the same regulatory elements as Epi TFs, a phenomenon we also validated in blastocysts. We propose that the ability of PrE and Epi-specific TFs to extensively bind and regulate the same gene networks contributes to ICM plasticity and allows rapid cell lineage specification by coordinating both activation and repression of divergent transcriptional programs.

---

Recent single-cell (sc) studies have highlighted the prevalence of plastic states characterized by coexpression of divergent lineage-determining TFs during cell specification (Farrell et al., 2018; Mittnenzweig et al., 2021). The mouse second cell fate decision is an in vivo paradigm to understand how lineage-specific TFs regulate differentiation of diverse cell types from common progenitors. In mice, Epi and PrE arise from a common progenitor in the early blastocyst, the inner cell mass (ICM). Around embryonic day 3.25 (E3.25), all ICM cells uniformly express the Epi TFs, SOX2, OCT4, and NANOG, along with the PrE TF GATA6 (Dietrich and Hiiragi, 2007; Plusa et al., 2008). By E3.5, the ICM loses its uniformity and starts exhibiting an intermingled distribution of cells expressing either NANOG or GATA6 which are predisposed towards the Epi or PrE fates, respectively (Chazaud et al., 2006; Plusa et al., 2008; Saiz et al., 2016b; Schrode et al., 2014; Xenopoulos et al., 2015). By E4.5, both lineages are established, specified PrE cells segregate to form an epithelial layer encapsulating the pluripotent Epiblast, and the blastocyst implants in the uterus, culminating preimplantation development (Gerbe et al., 2008; Plusa et al., 2008; Saiz et al., 2013).

ICM cells preferentially expressing GATA6, and subsequently its downstream targets SOX17 and GATA4, activate a transcriptional network promoting commitment to the PrE fate (Artus et al., 2011; Bessonnard et al., 2014; Koutsourakis et al., 1999; Morrisey et al., 1998; Schrode et al., 2014; Shimoda et al., 2007; Soudais et al., 1995). However, PrE precursors in early blastocysts can still switch to an Epi fate if the ratio of Epi/PrE compartments is affected (Saiz et al., 2020; Yamanaka et al., 2010), indicating that cells at this stage are plastic and not fully committed. This adaptability is lost in the late blastocyst (E4.5) suggesting that Epi and PrE fates are irreversibly committed at this stage (Grabarek et al., 2012; Yamanaka et al., 2010). Although Epi- and PrE-specific genes have been described using scRNA-seq in developing blastocysts (Mohammed et al., 2017; Nowotschin et al., 2019; Ohnishi et al., 2014), we lack understanding of how plasticity is maintained in the ICM and then quickly lost as cells are specified into two distinct lineages.

Embryonic stem (ES) cells cultured in vitro do not express GATA6 or any other PrE-specific genes and have been used to study regulatory mechanisms controlling the Epi pluripotency network (Boyer et al., 2005; De Los Angeles et al., 2015; Di Giammartino and Apostolou, 2016; Heurtier et al., 2019; Young, 2011). Similarly, extra-embryonic endoderm (XEN) cells derived from blastocysts have been used to characterize the PrE fate in vitro. However, XEN cells are a heterogenous population, more closely resembling the PrE post-implantation derivatives, parietal and visceral endoderm (Cho et al., 2012; Kunath et al., 2005). Alternatively, ectopic expression of PrE TFs like GATA6, GATA4 or SOX17 in ES cells, provide efficient temporally controllable systems, which have helped identify genes and signaling pathways driving PrE differentiation and how they are induced by these PrE TFs (Fujikura et al., 2002; Morris et al., 2010; Niakan et al., 2010; Schroter et al., 2015; Shimosato et al., 2007; Wamaitha et al., 2015).

To characterize the plastic state of bipotent ICM cells we used ES-cells carrying an inducible GATA6 transgene and profiled them immediately following GATA6 induction (2h). While GATA6 represses NANOG and other Epi TFs quickly, at this early differentiation timepoint both GATA6 and NANOG are hihgly expressed. We found that, during this stage of coexpression, GATA6 binds its motifs in cis-regulatory elements (CREs) of PrE genes and works as a pioneer TF to make them accessible for transcriptional activation. Simultaneously, GATA6 bound at an unexpectedly large fraction of CREs controlling Epi genes, leading to co-occupancy with the core pluripotency TFs, NANOG and SOX2. Surprisingly, we observed that transient binding of GATA6 to Epi CREs was followed by eviction of NANOG and SOX2 and redirection to GATA6-bound PrE CREs possibly contributing to robust activation of the PrE network while indirectly repressing Epi genes. The ability of GATA6 to bind at both Epi and PrE genes was evident also in uncommitted ICM cells in blastocysts. We propose that binding of NANOG and GATA6 to the same regulatory elements confers plasticity in ICM cells and allows rapid bifurcation into divergent lineages during blastocyst development.

## RESULTS

### GATA6 initiates rapid transcriptional and chromatin changes to induce PrE-like cells in vitro

To characterize how GATA6 and NANOG coregulate a plastic state and how GATA6 initiates PrE differentiation while inhibiting the alternative Epi fate, we employed doxycycline (Dox)-induced GATA6 expression in ES cells (Wamaitha et al., 2015). GATA6 expression was induced for 12h and cells cultured up to 96h **(Fig 1A)**. To identify the differentiation time points that resembled in vivo stages more closely, we used a scRNA-seq dataset (Nowotschin et al., 2019) generated from E3.5 and E4.5 blastocysts and compared them to bulk RNA-seq of GATA6-induced cells at each time point (**Fig 1B**). To achieve this, we first obtained normalized expression data using standard procedures specific to each data source (see Methods for details). Next, we fit a linear model to account for technical variability of the different platforms and performed principal component analysis (PCA) on the residuals to compare bulk and scRNA-seq timepoints. Reassuringly, trophectoderm (TE) cells, which are specified independently in the first cell fate decision, clustered away from all other data points. Each time point between 8 to 48h after Dox induction, transcriptionally clustered with a distinct cell type in blastocysts. At 8h, induced cells resembled ICM and Epi cells, while at 16h they clustered with PrE precursors present in early blastocysts (E3.5). Between 24 and 48h, in vitro cells resembled the PrE cells found at E4.5, suggesting that the process of PrE fate-determination had been completed. The 96h time point clustered away from the blastocyst cells suggesting that by this time point cells most likely resembled PrE derivatives, deeming them less relevant to study PrE specification mechanisms. FACS and immunofluorescence confirmed that, as reported (Wamaitha et al., 2015), almost all cells induce PrE targets and repress Epi genes by 48h (**Fig S1A, S1B)**. We then looked at changes in expression of genes forming the Epi and PrE transcriptional networks. Epi- and PrE-specific genes were defined by identifying genes differentially expressed in the two ICM-derived lineages. For higher stringency, only genes identified as Epi or PrE-specific in a second dataset (Mohammed et al., 2017) were included in downstream analyses. Genes more highly expressed in the PrE population, were included in the PrE transcriptional network (579 genes), and vice versa for Epi genes (221 genes). As shown in **Fig 1C**, the transcriptome progressively changed during differentiation, with a decrease of Epi-, and an increase of PrE-transcript levels. Differential gene expression analyses of the entire transcriptome also revealed progressive changes in expression, starting as early as 2h post GATA6 induction, with 639 genes silenced, and 920 activated (**Fig S1C, S1D**). In summary, these data show that within 48 hours, GATA6-induced ES cells approach a transcriptional state that resembles PrE-specified cells at E4.5.

**Figure 1.**
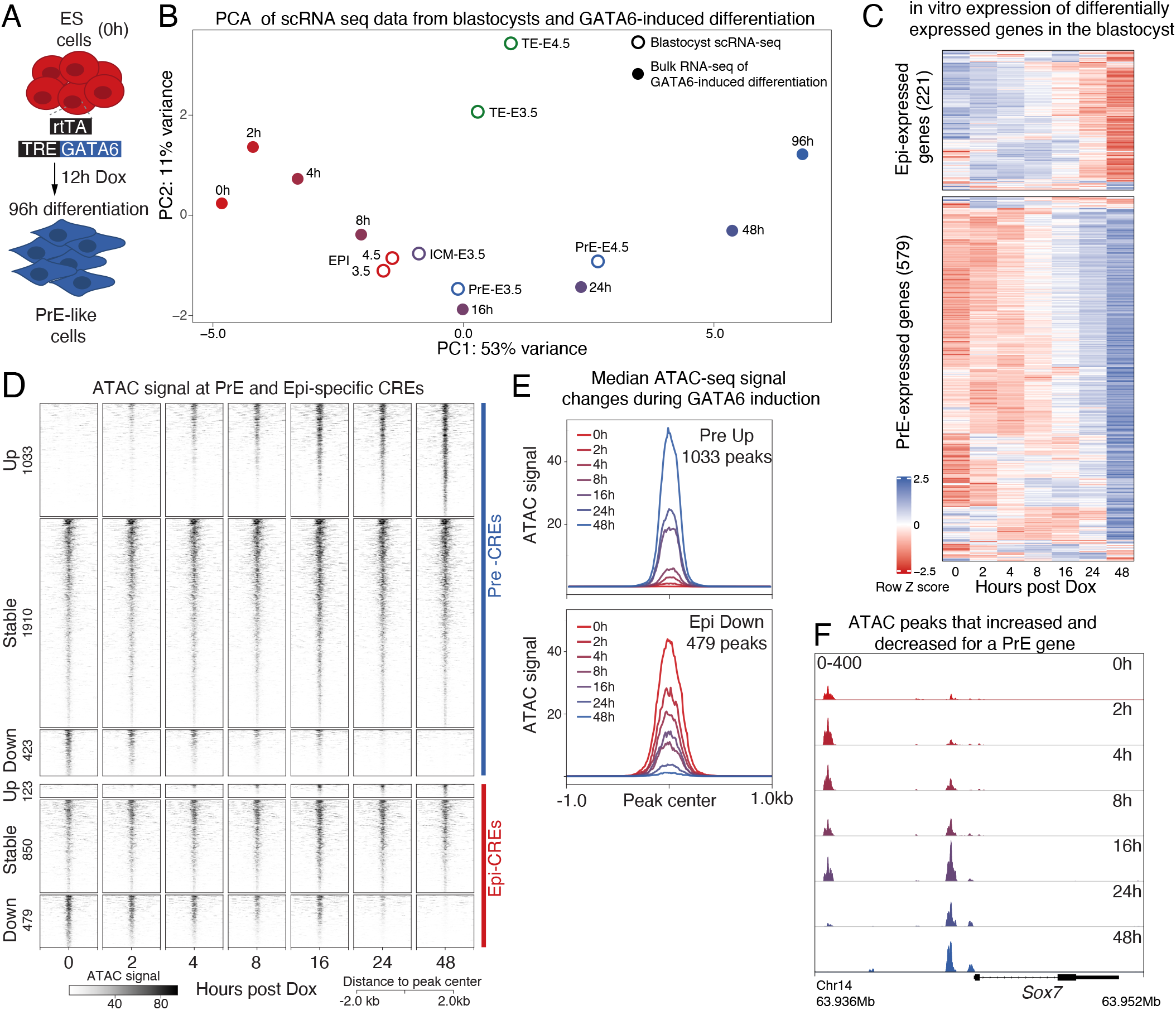
GATA6-driven in vitro differentiation recapitulates PrE specification in blastocysts and initiates rapid transcriptonal changes and chromatin remodelling. **A** Scheme depicting GATA6-driven in vitro differentiation of mES cells used to study PrE specification. **B** PCA of bulk-RNA-seq at several differentiation time-points (solid circles) and single-cell RNA-seq profiles of E3.5 and E4.5 blastocysts (open circles). The PrE precursor state was achieved by 16 hours of GATA6-driven differentiation and fully specified PrE-cells appeared at 24-48hours. **C** Heatmap of transcript z-scores of Epi- and PrE-specific genes shows that most blastocyst lineage-specific genes showed similar expression patterns during GATA6-driven differentiation. Epi and PrE-specific genes were defined using blastocyst scRNA-seq data. **D** Changes in chromatin accessibility measured by ATAC-seq at PrE and Epi CREs were quickly initiated within 2 hours of differentiation. Accessible regions at 48 hours were compared to 0 hours and classified as up, stable or down. **E** Median ATAC signal plots depict an increase in ATAC signal at 1033 PrE-CREs and decrease at 479 Epi-Cres. **F** Browser view at the Sox7 locus showing a progressive loss in ATAC signal at distal CRE and a gain in accessbility at a proximal CRE. This is an example showing that not all PrE CREs gained accessibility during PrE-specification. Loss of chromatin binding by transcriptional repressors can explain why some PrE-CREs lost accessibility

To probe into the dynamics of chromatin remodeling accompanying PrE differentiation, we profiled changes in chromatin accessibility using ATAC-seq (Buenrostro et al., 2015). This revealed that by 2h of GATA6 expression, changes in chromatin accessibility were more striking than those observed at the transcriptome level (**Fig S1C, S1D, S1E, S1F**), indicating that remodeling of chromatin landscape occurs at a faster pace than transcriptional changes. ATAC peaks were classified as Epi- or PrE-specific CREs using the closest TSS for a distance up to 50kb, a method that is widely used to annotate GWAS loci (Nasser et al., 2021). In both cases, the majority of CREs (1910) did not change accessibility during differentiation (**Fig 1D**). Although most PrE CREs that showed changes were associated with an increase in accessibility (1033), some showed a reduction (423). Conversely, most dynamic Epi CREs lost accessibility (479) while a few showed a gain (123). Gain of ATAC signal at PrE CREs was evident as early as 2h with a surge at 16h (**Fig 1E**, top). At Epi CREs, loss of accessibility also began at 2h and decreased gradually during differentiation (**Fig 1E**, bottom). The PrE 16h surge in accessibility is exemplified at the promoter-proximal CRE controlling *Sox7*, a PrE-specific TF activated by GATA6 (**Fig 1F**). In contrast, a distal *Sox7* CRE lost chromatin accessibility, possibly due to a change in binding of a *Sox7* repressor. This is a good example of why not all PrE CREs gain accessibility as it may have been expected. Similar chromatin remodeling dynamics were observed at ATAC peaks identified throughout the genome, without restricting analysis to Epi or PrE-specific CREs (**S1F, S1G**). These data demonstrate that GATA6-driven chromatin remodeling is initiated immediately following induction and that characterization of early events is crucial to understand how a single transcription factor initiates PrE differentiation while inhibiting the Epi transcriptional program.

### GATA6 functions as a pioneer TF to rapidly activate the PrE transcriptional network

To identify regulatory mechanisms contributing to changes in chromatin accessibility during differentiation, we mapped TF binding using CUT&RUN (Skene and Henikoff, 2017) and the distribution of histone tail modifications associated with active and repressed transcriptional states using CUT&TAG (Kaya-Okur et al., 2019). We detected endogenous *Gata6* mRNA, as early as 2h after addition of Dox, at levels comparable to those of the transgene (**Fig 2A**) showing that GATA6 quickly autoregulates its own expression. Other known downstream GATA6 targets such as *Gata4, Sox17, Hnf1b* and *Pdgfra*, showed increased transcript levels only by 4-8h (**Fig 2A**). To understand how GATA6 and its downstream targets regulate the PrE program, we mapped their binding during the course of differentiation. We first identified genome wide GATA6 peaks at early (2,4,8h), and late (48h) stages of differentiation and by comparing with ATAC data, saw that GATA6 binding could be divided into three peak-types of similar proportions: early binding at regions that were already accessible before GATA6 induction, early binding at regions that only become accessible after induction, and regions bound only at late differentiation stages (48h). These three GATA6-bound peak types showed an increase in accessibility following binding and were detected both at PrE-specific CREs (**Fig 2B, 2C**), and genome wide (**Fig S2A, S2B**). As defined for ATAC, GATA6 peaks were classified as PrE CREs if the closest TSS found within 50kb of the peak was a PrE-specific gene. Deposition of H3K4me3, a mark of active transcription, increased at promoter-proximal PrE-CREs (within 5kb) (**Fig 2B, S2C**), while H3K27ac, a mark of active enhancers, was deposited at promoter-proximal and -distal PrE-CREs (**Fig 2B, 2C)**. Irrespective of the timing of GATA6 binding (early or late), H3K27ac deposition trailed the gain of accessibility (**Fig 2B, 2C, 2D, S2A**) by a few hours likely reflecting the time required to recruit histone acetylation complexes following GATA6 binding. Together, this shows that PrE CREs bound by GATA6 are marked for activation leading to transcription of PrE-specific genes. Surprisingly, most CREs activated by GATA6 binding did not show enrichment of repressive histone marks (H3K27me3 and H3K9me3) at 0h, in undifferentiated ES cells (**Fig 2B, 2C**, example of a CRE with H3K27me3 at 0h shown in **S2B**). We propose that this contributes to the fast activation seen during PrE lineage commitment upon GATA6 induction, as these regulatory elements would not require removal of heterochromatin marks for activation.

**Figure 2.**
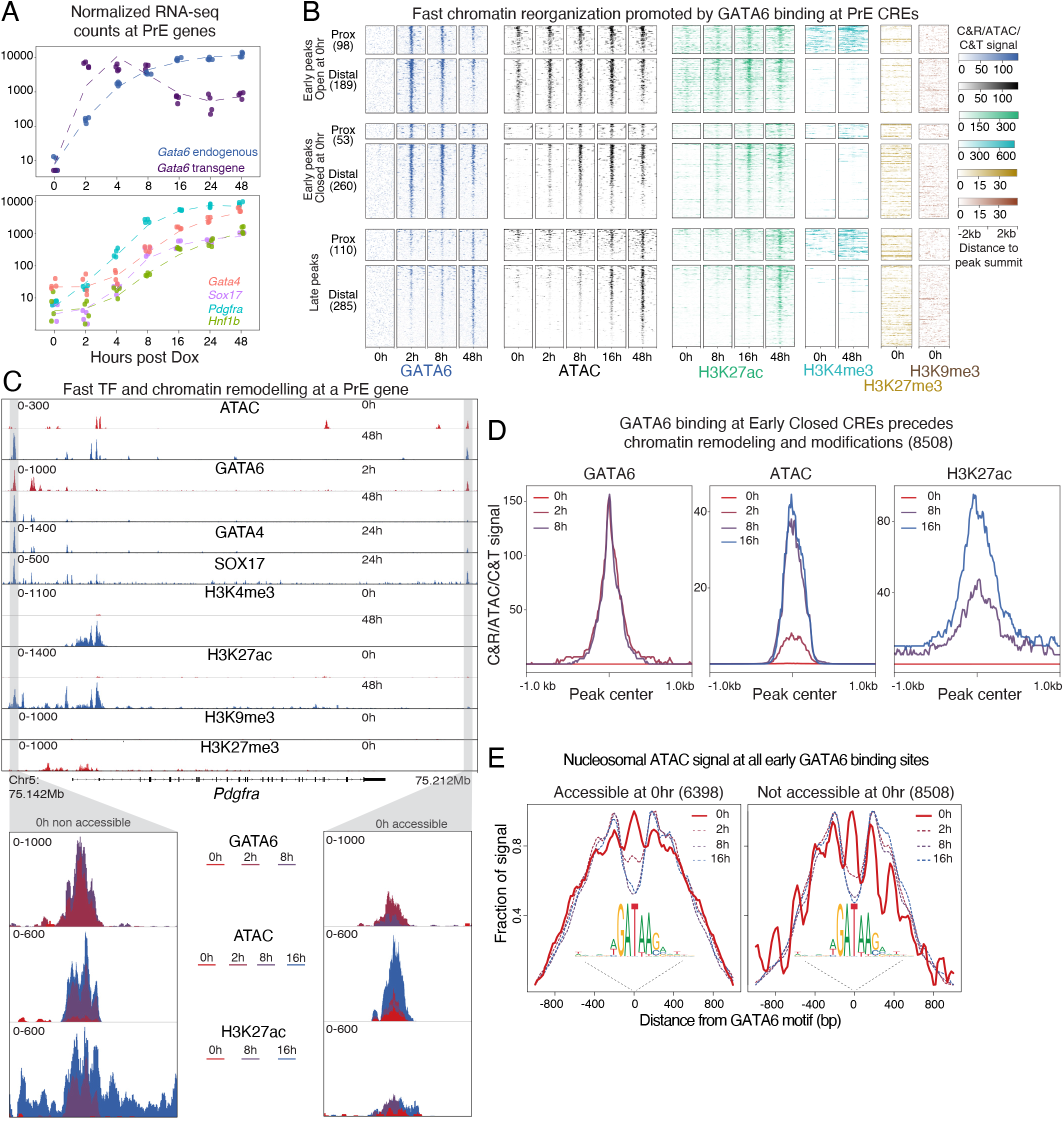
GATA6 binding at PrE-specific CREs precedes nucleosomal repositioning, changes in chromatin modifications and transcriptional activation of the PrE transcriptional program. **A** Normalized RNA-seq counts show a gradual increase in endogenous Gata6 levels that followed Dox-induced expression of the Gata6 transgene (top panel). Transgene and endogenous transcripts accumulated at similar levels. Gradual gain in transcript levels of key PrE TFs that are downstream targets of GATA6 is shown in the bottom panel highlighting transcript accumulation within 4 hours of GATA6 expression. **B** GATA6 peaks associated with PrE genes identified by CUT&RUN were classified as proximal (within 5kb of TSSs) or distal CREs and as late peaks (bound by GATA6 only at 48 hours) and early peaks (bound by GATA6 by 8 hours). Based on ATAC-seq signal, early peaks were further categorized into Closed at 0 hours or Open at 0hours. Heatmap compares differentiation-induced changes in GATA6 binding, corresponding ATAC-seq signal, as well as active (H3K27ac, H3K4me3) and repressive (H3K9me3, H3K27me3) histone marks at the different peak categories. **C** Browser view of the *Pdgfra* locus. Regions highlighted in grey depict changes in accessibility, binding by PrE TFs (GATA6, GATA4, SOX17), and changes in active and repressive histone marks. Two *Pdgfra*-putative CREs are shown, one accessible and one closed at 0 hours.**D** Median profile plots of normalized GATA6, ATAC, and H3K27ac signals at early GATA6 peaks closed at 0 hours shows that GATA6 binding preceded increases in accessibility and H3K27ac. **E** Nucleosomal fraction of ATAC-seq signal as determined by ATACseqQC at the indicated timepoints over GATA6 motifs in two types of GATA6 early peaks: accessible before the onset of differentiation (left panel), and accessible only after GATA6 binding (right panel). In both peak types nucleosomal positioning over GATA6 motifs dropped quickly following GATA6 induction with stronger decrease at sites that were not accessible at 0h.

While Dox induction led to immediate GATA6 binding to several CREs, others were bound only at 48h (**Fig 2B, S2A**, cluster 3). We hypothesized that late GATA6 sites require additional factors, only expressed later during differentiation, to gain complete chromatin accessibility. Therefore, we assessed binding of GATA4 and SOX17 at late-differentiation stages. As expected, there was a large overlap in binding targets for the three PrE TFs that was more prominent at the GATA6 sites bound only at 48h (**Fig S2D**). In addition, we ran HOMER motif enrichment at late GATA6 peaks by comparing with GATA6 early binding sites. The motif recognized by HNF1B, a TF shown to be important for visceral endoderm specification (Barbacci et al., 1999), was the most enriched motif (**S2D**). Notably, ATAC-seq footprint protection, which measures the likelihood of TF binding, showed mild protection of HNF1B motifs at 24h which increased by 48h, coinciding with GATA6 binding at its late target sites (**Fig S2E**). This suggests that binding of HNF1B aids GATA6 binding at late sites, along with GATA4 and SOX17.

To address if GATA6, like other GATA-family members (Mayran and Drouin, 2018; Tanaka et al., 2020), functions as a pioneer TF capable of accessing its motifs even if occluded by nucleosomes, we filtered ATAC-seq data at each time point to only include fragments with length longer than a nucleosome (>150 bp). We then plotted this filtered ATAC-seq signal at GATA6 motifs within early GATA6 peaks already accessible in undifferentiated cells (**Fig 2E**, left panel), and early GATA6 peaks inaccessible in undifferentiated cells (**Fig 2E**, right panel). In both instances, GATA6 motifs were occupied by nucleosomes in undifferentiated cells and were repositioned as early as 2h post induction. This supports a role for GATA6 as a pioneer TF in activating the PrE gene regulatory network. As GATA6 affects nucleosome occupancy within 2h of induction, ICM cells, which express GATA6, likely have most of their PrE CREs already bound by GATA6 poising them for rapid differentiation into the PrE fate.

### Inactivation of Epi CREs is preceded by transient GATA6 binding

In addition to activating the PrE transcriptional network, establishment of the PrE lineage requires repression of the alternative Epi fate. At the transcript level, the core Epi TFs, *Nanog* and *Sox2* were silenced progressively as PrE differentiation proceeded (**Fig 3A**). As shown previously (Mansour and Hanna, 2013), levels of the pluripotency factor OCT4 remained stable *(Pou5f1*, green). Since the protein levels of NANOG, the Epi-determining TF, were completely depleted by 12h (**Fig S3A**), we investigated how its genome-wide distribution changed during GATA6-induced PrE differentiation only at early time points (0 to 8h). As NANOG and SOX2 are known to bind and regulate self-renewal and pluripotency genes in ES cells (Boyer et al., 2005), we also investigated how SOX2 genome-wide distribution changed during GATA6-induced PrE differentiation.

**Figure 3.**
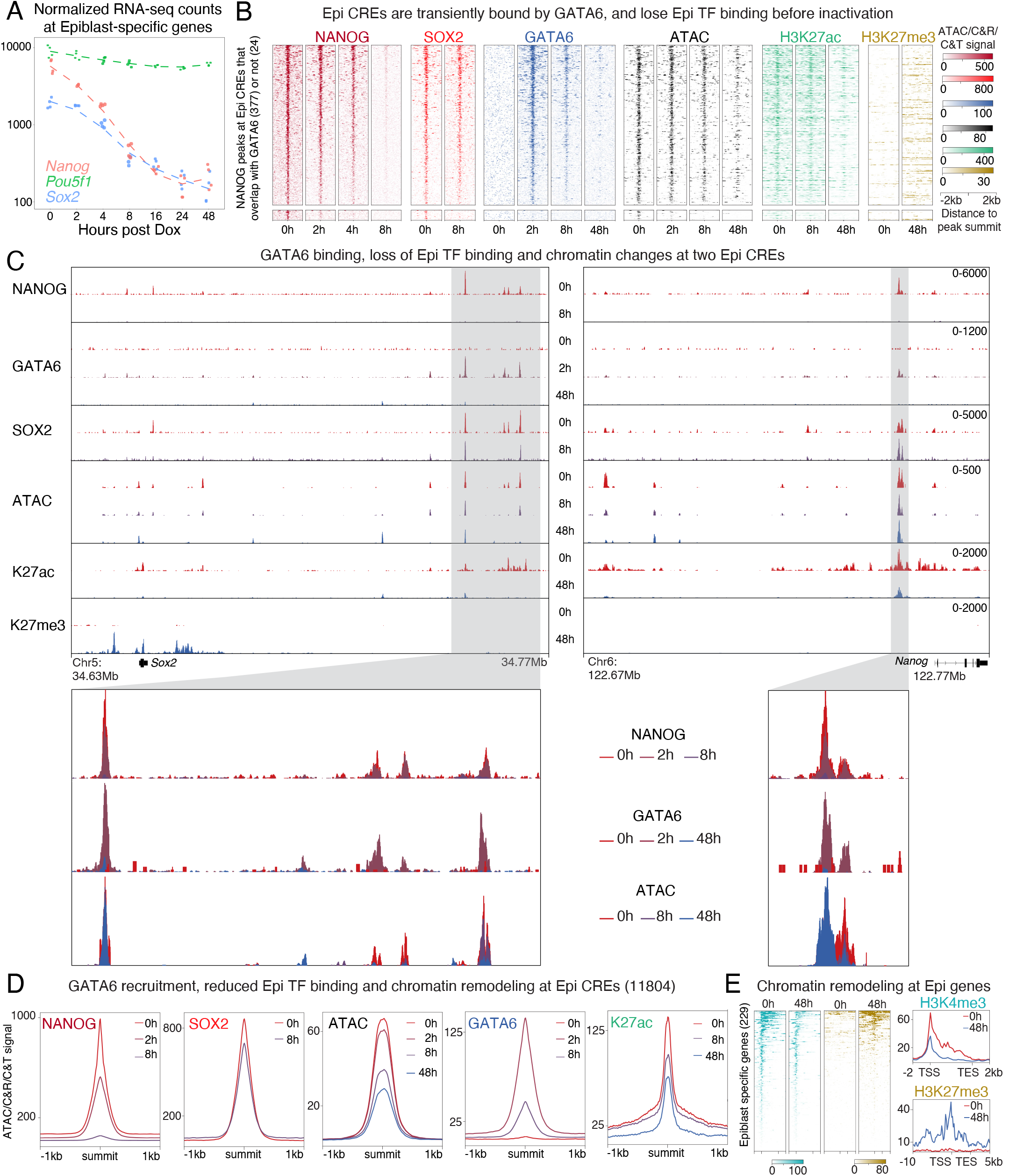
Inactivation of Epi CREs is preceded by transient GATA6 binding. **A** Changes in normalized RNA-seq counts of three Epi genes during differentiation. As in blastocysts, *Nanog* mRNA levels decrease faster rates than *Sox2*, while *Pou5f1* is less affected during differentiation **B** Heatmap shows gradual loss of NANOG and SOX2 at Epi CREs, concomitant with the transient binding of GATA6 at these loci, and progressive loss of ATAC signal and H3K27ac. A few CREs displayed increase in H3K27me3. **C** Browser view showing changes of NANOG, SOX2, and GATA6 binding, along with changes in ATAC signal, H3K4me3 and H3K27me3 at the *Nanog* and *Sox2* locus. While Nanog was silenced independently of H3K27me3, *Sox2* accumulated this mark over its gene bodies and surrounding regions. **D** Median profile plots depicting quick loss of NANOG and SOX2 with simultaneous GATA6 recruitment, and consequently a decrease in ATAC signal and H3K27ac. **E** Change in H3K4me3 and H3K27me3 surrounding Epi TSSs at 0 and 48 hour time points. Some Epi genes accumulated H3K27me3 during differentiation

At NANOG-bound regions, genome-wide and specifically at Epi CREs, we also detected strong SOX2 binding (**Fig 3B, 3C, S3B**). Strikingly, most of these sites were occupied by GATA6 starting as early as 2h post Dox induction (**Fig 3B**, top cluster, 377 peaks) and very few NANOG-SOX2 peaks were not bound by GATA6 (**Fig 3B**, bottom cluster, 24 peaks). In line with its role in repression of *Nanog* and other Epi genes, GATA6 was previously shown to bind near the promoter of some pluripotency genes (Wamaitha et al., 2015). By profiling the time points immediately following GATA6 induction, we show here that GATA6 binding at Epi CREs is much more extensive than previously described and likely impacts the Epi program more profoundly than previously thought. Contrary to PrE CREs, where GATA6 was bound throughout differentiation, Epi CREs and other NANOG-bound sites accumulated GATA6 transiently and only until 8h (**Fig 3B, 3C, 3D**-fourth panel). Concomitant with the transient GATA6 binding, NANOG diminished progressively from 2h onwards (**Fig 3B,D**, first panel). SOX2 binding at epi CREs diminished only slightly (**Fig 3B,D**, second panel). NANOG was completely displaced from its target sites by 8h which coincided with the progressive loss of chromatin accessibility (**Fig 3B**, black; **Fig 3D**, third panel, **Fig S3B**) and reduction in H3K27ac levels (**Fig 3B**, green; **Fig 3D**, fifth panel, **Fig S3B**). A loss of H3K4me3 was seen at the promoters of Epi genes (**Fig 3E**, blue) with a gain of H3K27me3 at some, but not all Epi promoters (**Fig 3E**, yellow). In contrast to activation of PrE CREs, where nucleosome repositioning preceded CRE activation, nucleosome occupancy at NANOG motifs within its target CREs, remained unchanged despite the reduction in accessibility and H3K27ac levels (**Fig S3C**).

Because GATA6 binding precedes the chromatin landscape changes occurring at NANOG-bound CREs, we speculate that GATA6 induces eviction of the Epi TFs, either directly or indirectly, to achieve repression of the Epi transcriptional network. Transient recruitment of GATA6 was seen at CREs regulating both *Sox2* and *Nanog* (**Fig 3C**) (Blinka et al., 2016; Zhou et al., 2014). These CREs showed a loss of NANOG-SOX2 binding, followed by reduction in chromatin accessibility which trailed GATA6 binding at 2h. Interestingly, *Sox2* accumulated H3K27me3, while *Nanog* only showed a decrease in H3K27ac with no accumulation of H3K27me3 (**Fig 3C**). Since *Nanog* levels decreased faster than *Sox2* during PrE differentiation, we propose that the repressive mechanisms employed by GATA6 at the two genes may reflect the difference in rate of transcriptional silencing.

We wanted to further understand if GATA6 induces silencing of the Epi program by direct or indirect binding. Since NANOG- and SOX2-bound regions are highly accessible (**Fig 3B, S3B**), it is possible that GATA6 is attracted to these sites simply because of their high accessibility. Additionally, if the detected GATA6 peaks at NANOG-bound regions contained the GATA6 recognition motif, it would argue that GATA6 binding could be direct. Indeed, we found GATA6 motifs within CREs that were commonly bound by both NANOG and GATA6 at 2h suggesting that inactivation of these regions is mediated by direct GATA6 recruitment (**Fig S3D**). To then address why NANOG-bound CREs recruited GATA6 less stably than PrE CREs, we compared the density of GATA6 binding motifs within the two types of GATA6 targets. Regions bound by NANOG and GATA6 contained lower density of GATA6 motifs as compared to GATA6 peaks not overlapping with NANOG at 0h (**Fig S3D**, boxplot). These peaks, which we consider to be non-PrE GATA6 targets, also contained GATA6 motifs of weaker strength as measured by sequence similarity to the canonical GATA6 motif (**Fig S3D**, violin-plot). Analysis of CUT&RUN cut-site probability at GATA6 motifs located both in NANOG 0h overlapping (Epi specific) and non-overlapping (PrE-specific) peaks provided further support that GATA6 does indeed occupy its motifs directly at both peak types **(Fig S3E**). This difference in GATA6 motif strength and density may explain why Epi CREs bind GATA6 only transiently while PrE CREs bind GATA6 more stably. It is also likely that GATA6 binding at Epi CREs is facilitated by NANOG occupying these regions and maintaining a highly accessible state. In line with this, GATA6 binding is reduced at 8h, when NANOG levels decrease, together with lower accessibility.

### Evicted pluripotency factors transiently occupy GATA6-bound PrE genes

Despite being displaced from Epi CREs, NANOG and SOX2 protein levels remained stable for at least 4h post Dox treatment (**Fig S3A**). Unexpectedly, we detected a transient doubling in the total number of NANOG and SOX2 peaks at 2h compared to 0h, which decreased back to initial numbers by 4h (total peaks identified for NANOG: 0h-19995, 2h-41877, 4h-23519) (**Table S1**). Since NANOG/SOX2-bound CREs were capable of recruiting GATA6, we wondered if the NANOG/SOX2 peaks gained at 2h were associated with GATA6-bound CREs. Strikingly, a large number of 2h-specific NANOG peaks appeared at GATA6-bound CREs made accessible after GATA6 binding at 2h (**Fig 4A**, blue cluster). Interestingly, while peaks that were bound by NANOG before differentiation began, progressively lost NANOG (**Fig 4A**, red cluster), sites made accessible by GATA6 (blue cluster) showed increased NANOG binding (**Fig 4B, 4C, 4D**). Importantly, GATA6-bound PrE CREs also accumulated NANOG and SOX2 (**Fig S4A**). These observations led us to conclude that NANOG and SOX2 evicted from Epi sites were redirected to GATA6-bound CREs. To validate this observation with a different technique, we performed chromatin immunoprecipitation coupled with sequencing (ChIP-seq) for NANOG at 0 and 2hrs. In agreement, with our observations from CUT&RUN, the transient increase in NANOG binding at GATA6-bound CREs, specifically at 2h, was also evident by ChIP-seq (**Fig S4B**).

**Figure 4.**
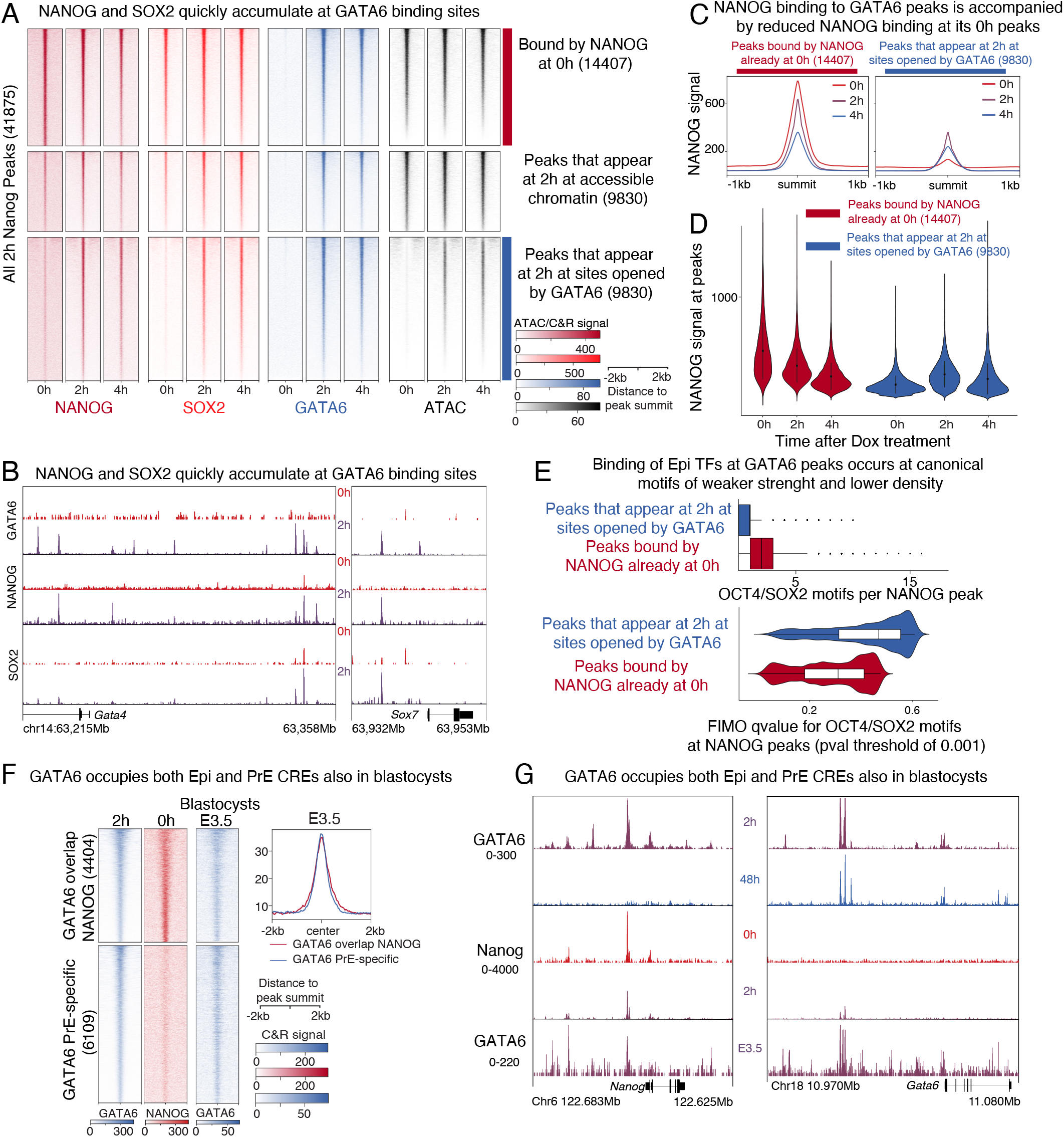
Evicted pluripotency factors transiently occupy GATA6-bound PrE regions. **A** Heatmap showing NANOG, SOX2, and GATA6 binding, together with ATAC signal at regions of the genome bound by NANOG at 2 hours. These NANOG peaks were divided in three types: occupied already at 0h (cluster 1-red), bound de novo by NANOG at 2hrs and accessible at 0hr (cluster 2), and bound de novo by NANOG at sites closed at 0h (cluster 3 -blue). **B** Representative browser view showing the de novo recruitment of NANOG and SOX2 at 2 hours at GATA6 peaks found at putative CREs controlling PrE genes GATA4 and SOX7.**C** Median profile plots showing a reduction in NANOG binding at NANOG 0h peaks (left panel, cluster 1 from A) with a simultaneous increase in NANOG binding at peaks opened by GATA6 binding (right panel, cluster 3 from A). **D** NANOG CUT&RUN signal from 0, 2, and 4 hours post Dox induction plotted at peaks comprising clusters 1 and 3 in 4A. The violin plot shows a decrease of read density at cluster 1 peaks with a simultaneous increase at cluster 3 peaks. **E** Peak per motif density and affinity of OCT4-SOX2 binding motifs within NANOG peaks that appear at 2 hours (blue) compared to NANOG peaks defined at 0 hours (red). **F** Extensive binding of GATA6 to Epi-CREs also occurs in blastocysts. Heatmap and median profile plot comparing GATA6 Cut&RUN binding in E3.5 blastocysts, to in vitro GATA6 and NANOG binding at 0h and 2h of GATA6-driven differentiation. **G** Example of GATA6 binding both at Epi *(Nanog)* and PrE loci *(Gata6)* in blastocysts.

We next considered the possibility that NANOG and SOX2 were redirected to GATA6-bound regions at 2h because these sites contain recognition motifs for the pluripotency factors. To address this, we focused on GATA6 target sites to look for enrichment of the murine OCT4/SOX2 consensus motif, which is better defined than the NANOG motif. We found that GATA6 peaks indeed contained the OCT4/SOX2 motif, although at lower density (**Fig 4E**, boxplot) and of weaker binding affinity (**Fig 4E**, violin-plot) than the motifs within pluripotency CREs (peaks bound by NANOG-SOX2 at 0h). The presence of binding motifs for Epi TFs, within GATA6-bound CREs, suggests that the PrE transcriptional network can be directly regulated by pluripotency TFs. Together with **Fig 3**, these data point towards a mechanism whereby GATA6 and NANOG regulate common CREs which may allow them to control lineage specification into either the PrE or Epi in blastocysts.

To address if GATA6 binds Epi and PrE CREs in unspecified ICM cells in vivo, we used CUT&RUN to profile GATA6 binding in early (E3.25-3.5) blastocysts (staged as in **Fig. S4C**), a stage when NANOG and GATA6 are coexpressed. Because specification of PrE or Epi fate is a rapid process, not occurring at the same time and same pace in all ICM cells, we suspected that in blastocysts, GATA6 binding at its target sites could be very transient. To reliably recover GATA6 targets in early and late blastocysts, we modified the CUT&RUN protocol to include light chromatin fixation. Additionally, instead of bead-assisted binding, we adapted methodology used for immunofluorescence in blastocysts (Plusa et al., 2008; Saiz et al., 2016a), which involves manual handling of embryos under the microscope at each step of the protocol and between washes. Our modified protocol prevented loss of biological material, preserved antibody recognition and physical attributes of the embryo **(Fig. S4C)**, and significantly improved our ability to recover high quality genome-wide TF binding data. In blastocysts, GATA6 binding was detected at regions bound in vitro by NANOG and GATA6 (**Fig 4F**, top cluster), and also at GATA6-specific peaks (**Fig 4F**, bottom cluster). Importantly, both GATA6 peak types exhibited similar GATA6 binding signal intensity (**Fig 4F**, top right plot). This demonstrates that the binding of GATA6 to both Epi and PrE CREs occurs in vitro and in blastocysts. A good example of GATA6-NANOG shared CREs is evident at both the *Nanog* and *Gata6* loci (**Fig 4G**). This data supports the idea that GATA6 indeed occupies and regulates the same targets as NANOG in developing embryos. Together, our in vitro and in vivo observations suggest that Epi and PrE TFs share common target sites within the Epi and PrE transcriptional networks. This extensive target sharing may facilitate rapid bifurcation of precursor cells into divergent lineages.

### Reshufling of TF binding is accompanied by rapid reorganization of local chromatin interactions

As PrE differentiation is associated with TF binding redistribution and changes in chromatin landscape we next investigated if these changes affected global threedimensional (3D) genome organization. Chromatin conformation assays have shown that large chromosomal domains, several megabases in length, can reside in either the active (A) or repressive (B) nuclear compartments depending on their transcriptional state and the histone modifications they harbor (Rocha et al., 2015). Because PrE differentiation was associated with changes in H3K27ac across the genome (**Fig 2B, 3B**), we first used Hi-C to profile 3D-genome organization of undifferentiated (0h) and PrEdifferentiated (48h) cells. Despite changes in chromatin landscape, genome organization at the compartment level was markedly similar in undifferentiated and differentiated cells. We performed PCA, and PC1 eigenvalue scores allowed identification of A and B compartment composition at 250kb bins. As an example, organization of compartments on chromosome 1 is depicted in **Fig 5A**, which shows how it remained largely unchanged even after 48h of differentiation.

**Figure 5.**
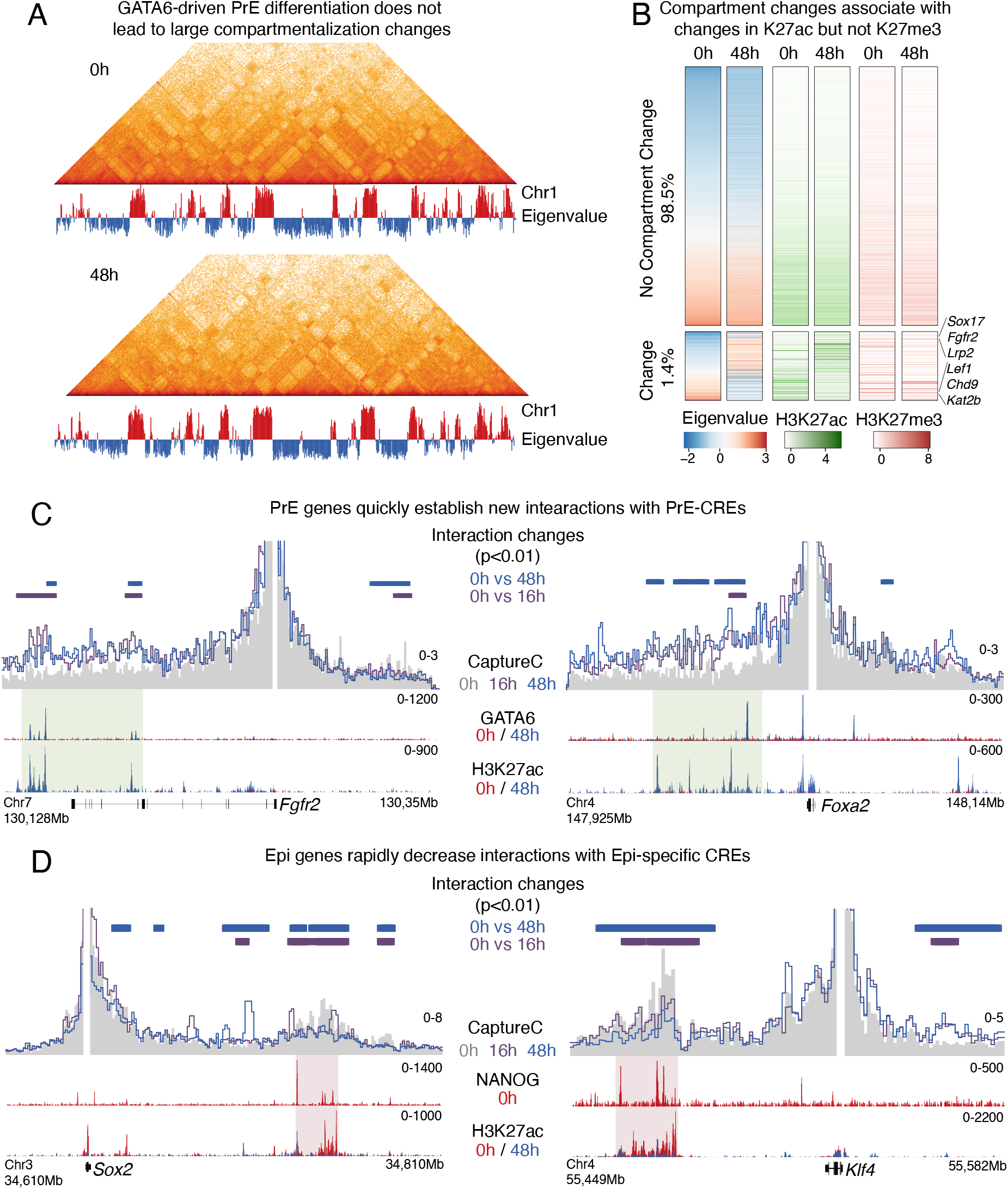
GATA6-induced chromatin changes are followed by fast rewiring of interactions between Epi and PrE-specific genes with their CREs. **A** Intra-chromosomal interactions over chromosome 1 detected by Hi-C and first principal component eigenvalue show that nuclear compartments composition was mostly unchanged during differentiation. **B** Heatmap showing eigenvalues of first principal component of Hi-C performed at 0 and 48 hours. Less than 2% of all 250kb-bins change their compartment during PrE differentiation (bottom cluster). Most 250kb-bins did not change compartment (top cluster). Clusters are not represented to scale. Changes in compartments were associated with changes in H3K27ac but no changes were observed for H3K27me3 levels. Bins that changed from compartment B to A gained K27ac signal and bins that changed from B to A, lost this histone modification. Bins were sorted based on eigenvalue at 0 hours. Examples of Epi and PrE-specific genes whose compartment localization changed during differentiation are shown. **C** and **D** In contrast to stable compartment status, interactions between gene promoters and their putative CREs were quickly remodeled within 16 hours of GATA6 expression. PrE genes shown in C, quickly increased interactions with sites bound by GATA6 that gained H3K27ac during differentiation. Epi genes in contrast, reduced interactions with CRES that were occupied by NANOG at 0h and that decrease H3K27ac following GATA6 expression. Capture-C data is shown as the average signal of two replicates using bins of 1kb. Green shaded area represents putative CREs with increased interactions with PrE-specific genes while red shaded areas highlight Epi-CREs that lose interactions with putative Epi-specific genes. P values were calculated using DESeq2 and comparing Capture-C signal of 2 replicates over overlapping 5kb windows across the regions shown in this figure. Comparisons were done between 0h and either 16h or 48h. Adjusted p values lower than 0.01 were considered as threshold for significance. Horizontal bars represent windows considered as statistically significantly different for these comparisons.

Genome-wide, just 1.4 percent of the entire genome showed differences in compartmentalization upon PrE differentiation (**Fig 5B**, bottom cluster). These genomic regions showed changes in H3K27ac levels and included PrE-specific genes like *Sox17, Fgfr2* and *Lrp2* which gained H3K27ac and were reorganized from inactive to active compartment. On the other hand, Epi genes such as *Lef1*, *Chd9*, and *Kat2b* lost H3K27ac and moved from the A to B compartment. Interestingly, genes such as Nanog, *Sox2*, *Gata6* and *Gata4*, that showed dramatic changes in transcript levels during differentiation, did not change compartments.

Besides large-scale chromatin reorganization, gene transcription can impact and be impacted by local interaction changes involving proximal and distal regulatory elements.

TFs have been shown to influence genome topology to drive differentiation-associated expression changes (Apostolou et al., 2013; Dall’Agnese et al., 2019; de Wit et al., 2013; Denholtz et al., 2013). While some reports have shown that changes in genome topology can be linked to transcriptional changes (Beagan et al., 2020; Di Giammartino et al., 2019; Freire-Pritchett et al., 2017; Rubin et al., 2017; Stadhouders et al., 2018), contrasting studies at developmentally regulated genes (Despang et al., 2019; Espinola et al., 2021; Gha-vi-Helm et al., 2014; Ing-Simmons et al., 2021; Williamson et al., 2019) have shown that transcriptional changes can occur independently of chromatin structure reorganization. These opposing results suggest that the contribution of 3D-genome interactions is locus and context specific. Since PrE specification involves rapid chromatin remodeling, we wondered if this short time frame was sufficient to alter genome topology at key PrE and Epi genes. To address this, we performed Capture-C to assess changes in interactions between CREs controlling Epi and PrE genes that showed the highest transcriptional changes between undifferentiated (0h) and differentiated stages (48h). At all PrE-specific genes tested, with the exception of *Fgf3*, we detected a gain in interaction between the promoter and putative CREs marked by gain of H3K27ac (**Fig 5C, S5**) within 16h of differentiation. Contrarily, most Epi-genes assayed (except for *Morel, Pou5f1*, and *Fgf4)*, were associated with a loss of interaction with their CREs, which lost H3K27ac during PrE differentiation (**Fig 5D, S5**). Interestingly, we could not identify distal lineage-specific CREs for any of the four genes without changes *(Fgf3, Fgf4, Morel, Pou5f1)* in interactions during differentiation. As these genes are likely regulated only by proximal CREs, this might explain the stability of 3D interactions upon GATA6 induction. Genes not expressed in either lineage were used as controls and as expected, showed no change in interactions from their promoter regions (**Fig S6**, *Cdx2* and *Pax5)*. In addition to increase in H3K27ac, all PrE CREs tested were associated with GATA6 binding in their vicinity. Likewise, Epi CREs where interactions decreased during differentiation, showed a loss of NANOG and SOX2 binding initiated by GATA6 binding at these sites. Thus, even though PrE differentiation is associated with limited large-scale genome reorganization, we observed significant changes in interactions between CREs controlling genes forming the Epi and PrE networks. Together our data show how GATA6, the PrE-specifying TF, initiates a cascade of interdependent mechanisms to regulate the genome in order to specify the PrE lineage and inhibit the alternative Epi fate. fox jumped over the lazy dog.

## DISCUSSION

Multicellular organisms comprise of diverse cell-types which often arise from a common progenitor. scRNAseq studies have highlighted how this is accomplished by combinatorial TF modules that promote multifurcation of cell-types from common multipotent progenitors (Mittnenzweig et al., 2021). Using GATA6-driven differentiation of ES cells, we propose how NANOG and GATA6 maintain plasticity in bipotent ICM cells, and how higher GATA6 levels preferentially promote a PrE fate. Cells between 2 to 8h of differentiation coexpress GATA6 and NANOG, and resemble the ICM in E3.25 blastocysts or uncommitted progenitor cells in later blastocysts. During co-expression (at 2h) we detected GATA6 binding along with the pluripotency TFs at both Epi and PrE CREs. This is corroborated by our observations in blastocysts, where we also detected GATA6 binding both at Epi and PrE CREs (**Fig 4**). In uncommitted ICM cells, when NANOG and GATA6 levels are equivalent, we propose that both factors are bound at their recognition motifs within Epi and PrE CREs, maintaining ICM cells in a poised state to adopt either cell fate.

When higher GATA6 levels are reached, relative to NANOG (4h in vitro), we observed that pluripotency TFs are evicted from Epi CREs and redirected to PrE CREs, which we propose aids GATA6 in achieving quick activation of the PrE network. Similar TF mobilization has been described during reprogramming of fibroblasts into induced pluripotency stem cells. The reprogramming TFs - OCT4, SOX2 and KLF4 (OSK) were shown to inactivate the somatic-cell transcriptional network by disengaging somatic TFs from their target sites and redistributing them to pluripotency loci engaged by OSK (Chronis et al., 2017) to facilitate activation of the pluripotency program. Similarly, the transition of naive to primed pluripotency in Epi cells of the implanted embryo is facilitated by redistribution of OCT4 (Buecker et al., 2014). In this case, OCT4 relocalization from naïve-to primed-pluripotency genes, was initiated by the primed-pluripotency specific TF, OTX2. OCT4 redistribution was preferentially seen at sites carrying abundant and strong OTX2 recognition motifs. Similar to these reported mechanisms, we propose that the inherent differences in the strength and density of GATA6 binding sites in PrE-compared to Epi CREs enables a shift in binding patterns of pluripotency factors when GATA6 levels are high, thus promoting an exit from plasticity and differentiation into PrE.

In support of this we found that PrE CREs contain higher density of high-affinity GATA6 recognition motifs compared to Epi CREs. This difference in motif affinity and density could provide a distinguishing feature enabling GATA6 to achieve contrasting changes at PrE and Epi CREs. A scenario can be envisioned where CREs carrying higher densities of strong TF motifs would be bound longer and more stably by the TF, marking them for transcriptional activation. Contrarily, less abundant, lower affinity binding sites could lead to only transient occupancy by the TF, not allowing sufficient time for activation, and instead marking them for inactivation by recruitment of histone deacetylases. A related observation was recently made where cells of the caudal epithelium that contain neural and mesodermal progenitors use binding of the same TF (SOX2) at sites of different affinity to elicit opposing regulatory responses in cells differentiating into divergent paths (Blassberg et al., 2020). In our study, we observed that CREs activated during PrE differentiation were bound by GATA6 starting at 2hrs and continued in most cases until 48hr (**Fig 2B**). CREs inactivated by GATA6, on the other hand, were bound by GATA6 only transiently at 2hr (**Fig 3B**), supporting that GATA6 residence time at its targets could affect transcriptional activity. This, however, remains to be tested thoroughly. Combinatorial binding by other lineage-restricted transcription factors could also contribute to marking loci for activation and to allow for input signal integration by different signaling pathways. This has been studied extensively and described in lineage specification of multiple cell types (De Val et al., 2008; Junion et al., 2012; Luna-Zurita et al., 2016; Spitz and Furlong, 2012; Stefflova et al., 2013). CREs controlling the PrE transcriptional network carry high affinity binding sites for GATA6 (**Fig S3D**) and are also bound by its downstream targets SOX17 and GATA4 (**Fig S2D**) potentially functioning to mark PrE genes for robust activation rather than repression.

In addition to NANOG and GATA6 levels, FGF/ERK signaling via FGF4 is essential for lineage fate choice in the ICM. In *Fgf4* ^*-/-*^ blastocysts, all ICM cells adopt an Epi fate at implantation, even though GATA6-expressing PrE-precursors are initially present, showing that PrE fate determination is dependent on FGF4/ERK signaling (Chazaud et al., 2006; Feldman et al., 1995; Grabarek et al., 2012; Kang et al., 2017; Kang et al., 2013; Molotkov et al., 2017; Nichols et al., 2009; Ohnishi et al., 2014; Saiz et al., 2020; Yamanaka et al., 2010). A mechanism where FGF4 regulates the ability of ES cells to exit pluripotency and differentiate into PrE was described recently (Hamilton et al., 2019). Using an in vitro system where levels of ERK activation can be modulated, Hamilton et al. showed that enhancer activity was proportionally affected merely by reversible dissociation of cofactors and the transcriptional machinery without TF redistribution. This provides an explanation for how stochastic fluctuations in FGF4 availability can be incorporated at the transcriptional level to promote divergent cell-fate determination in ICM cells. It is noteworthy that the extent of FGF/ERK activation used in this study, while sufficient to affect the degree of pluripotency was not enough to upregulate GATA6. This suggests that the observed changes in FGF signaling, recapitulated stochastic initiation of fate determination in early blastocysts where skewed ratios of GATA6 and NANOG are not yet established. Our study on the other hand, describes events starting from coexpression to establishment of skewed GATA6:NANOG ratio and hence recapitulates the process by which PrE fate-determination is initiated and completed.

The extensive TF redistribution evident during PrE specification is accompanied by rapid local chromatin reorganization. GATA6 binding at PrE and Epi CREs initiates a cascade of changes at multiple layers of chromatin. By altering the levels of H3K27ac, GATA6 regulates enhancer activity, which coincides with rewiring of genome structure. H3K27ac levels have been shown to influence enhancer-promoter interactions (Beagan et al., 2020; Rubin et al., 2017), however it is unclear if enhancer activity results in rewiring of interactions, or vice versa. It is plausible that in unspecified ICM cells, fluctuations in enhancer activity are first initiated and once GATA6 is expressed above a certain threshold compared to NANOG, PrE enhancers are stably activated (and Epi enhancers inactivated) leading to rewiring of enhancer-promoter interactions to facilitate continued enhancer activity. A temporal resolution of the two mechanisms, changes in histone modification and chromatin rewiring, would allow for the stepwise generation of a feedforward loop driving successive and unidirectional lineage commitment. Alternatively, it is also possible that TF-mediated multilayered chromatin regulation provides redundancy to ensure robust gene regulation. Further studies will be required to address these possibilities, and the blastocyst, owing to its simplicity and temporally distinct stages, provides an excellent self-contained system to gain mechanistic insights in vivo.

## METHODS

### Cell line, culture conditions, and in vitro differentiation

Previously generated mouse embryonic stem cells (mES) expressing a single copy a transgene containing a bidirectional TET-responsive element driving expression of Doxycycline-inducible GATA6-FLAG and DsRed2 were a kind gift from Dr. Niakan’s laboratory (Wamaitha et al., 2015). Cells were maintained at a density of 1 million cells per 10cm gelatinized plates at 37°C x% O2, x% CO2 in 2i media (50% DMEM/F12, 50% Neurobasal, Penicillin-Streptomycin, GlutaMAX, 2-Mercaptoethanol, N2, B27, 0.3nM PD0325901, 0.1 nM CHIR9902, and 2×10^5^units/mL LIF). Differentiation into primitive endoderm was induced in serum containing ES media (Knockout DMEM, 15% FBS,1% GlutaMAX, 1% Penicillin-Streptomycin, 1% 2-Mercaptoethanol, 1% Sodium Pyruvate, 1% MEM NEAA, 2×10^5^units/mL LIF) containing doxycycline (1ug/mL) for 2 to 12 hours. For time points greater than 12 hours, mES cells were induced in serum media containing doxycycline for 12 hours and switched to serum media in the absence of doxycycline for the remaining time. generated mouse embryonic stem cells.

### Fluorescence assisted cell sorting

Cells were harvested using 10mM EDTA made in PBS. After incubation in EDTA for 10 mins, cells were dislodged by pipetting to ensure a suspension of single cells, following which cells were spun and resuspended in 500μl of MACS buffer (PBS, 2% FBS, 1mM EDTA). Cells were double stained with preconjugated PDGFRA (FITC) and PECAM (APC) antibodies by adding 0.5μl of each antibody. Cells were incubated at 4°C for 20 mins following which they were washed twice in MACS buffer. Cell pellets were finally resuspended in 800μl MACS buffer and analyzed for proportions of FITC positive/APC negative cells.

### Immunofluorescence

Cells were seeded on 6-well dishes in 2i, at a density of 25,000 cells/well. After Dox induction, cells were rinsed twice in PBS and fixed using 4% formaldehyde for 10 mins, following which cells were washed thrice in PBS and then permeabilized for 10 mins at room temperature using 0.5% Triton X-100 made in PBS (0.5% PBST) supplemented with 100mM glycine. Cells were again washed thrice in PBS and incubated in 2% Horse serum, in a humidified chamber at 37°C for 30 mins. Cells were then incubated in primary antibodies (SOX17-R&D Systems, AF1924, SOX2-Millipore, ab5603) diluted at 1:500 in 2% horse serum, 0.5% Triton-X-100 made in PBS, at 4°C overnight. Cells were then washed thrice in 0.1% PBST and incubated in secondary antibodies (prepared in PBS at 1:1000) for 1 hour at room temperature, protected from light. Secondary antibody was then replaced with Hoechst solution (10μg/ml in PBS) for 2 mins and then cells were thoroughly washed in 0.1% PBST before imaging at 10X on an inverted epifluorescence microscope.

### Western blot

Whole cell extracts were prepared in 1X Radioimmunoassay buffer (RIPA, 50 mM Tris-Cl pH 8,150 mM NaCl, 2mM eDTA pH 8, 1% NP40, 0.5% sodium deoxycholate, 0.1% SDS) supplemented with protease inhibitor cocktail (Roche). Protein content was estimated using the Peirce BCA kit, and equal amounts of protein (20μg) resolved on 10% SDS-PAGE gels. Proteins were transferred onto PVDF membrane (Immobilon-FL), followed by blocking (5% milk made in TBST) and incubation with primary antibodies. Proteins were detected using HRP-conjugated secondary antibodies. Primary antibodies (and dilutions) are: Rabbit anti-NANOG (ab80892; 1:1000), Rabbit anti-SOX2 (ab92494; 1:1000), histone-H3 (ab176842; 1;1000).

### RNA extraction, RNA-seq library preparation and sequencing

RNA from four replicates at each time point was isolated using trizol. After confirming that the RNA integration number for each sample was above 8, libraries were prepared using TruSeq Stranded mRNA prep kit with PolyA purificaton and sequenced on HiSeq 2500 using SE50.

### ATAC-seq library preparation and sequencing

ATAC-seq was performed as described in (Corces et al., 2017). Briefly, cells at each time point were dissociated using Accutase, and 50,000 cells were subjected to the tagmentation reaction. Cells were first washed in resuspension buffer (10 mM Tris-HCl pH 8.0, 10 mM NaCl, and 3 mM MgCl2 in water), following which nuclei were isolated in 1 ml lysis buffer (10 mM Tris-HCl pH 8.0, 10 mM NaCl, 3 mM MgCl2, 0.1% NP-40, 0.1% Tween-20, and 0.01% Digitonin in water) on ice for three minutes. Nuclei were rinsed once in wash buffer (10 mM Tris-HCl pH 8.0, 10 mM NaCl, 3 mM MgCl2, and 0.1% Tween-20) and tagmentation was carried out using 2.5ul Tn5 transposase (Illumina 15027865) for exactly 30 mins. Following tagmentation, DNA was purified using the Zymo DNA Clean and Concentrator kit (Zymo, D4033). Libraries were prepared using Q5 polymerase and unique indices were added to each sample. First, gap filling was performed at 72°C for 5mins followed by five cycles of 98°C, 20secs, 63°C, 30secs, and 72°C 1min. After initial amplification, tubes were held on ice, while quantitative PCR was run on 1 μl of the pre-amplified library to determine additional number of cycles needed. Libraries were sequenced on HiSeq2500 using PE50.

### CUT&RUN

CUT&RUN was performed as previously described (Skene et al., 2018) with small modifications. Briefly, mES cells were dissociated using Accutase, counted, and 100,000 cells/replicate were pelleted at 600g for 3 minutes at room temperature. Supernatant was discarded, cells were resuspended in Wash Buffer (20 mM HEPES pH 7.5, 150 mM NaCl, 0.5 mM Spermidine, 1x Protease inhibitor cocktail), and pelleted at 600g for 3 minutes at room temperature. BioMag® Plus Concanavalin A beads (Bangs Laboratories) were equilibrated in Binding Buffer (20mM HEPES pH 7.5, 10mM KCl, 1mM CaCl_2_, 1mM MnCl_2_). Cells were resuspended in Wash Buffer, mixed with a slurry of equilibrated Concavalin A coated magnetic beads, and rotated for 10 minutes at room temperature. Per 100,000 cells, a 10μl bead slurry was used. Beads were placed on a magnetic separator and supernatant was discarded. Beads were resuspended in Wash Buffer containing 2mM EDTA, 0.1% bovine serum albumin, 0.05% Digitonin, and 1:50 dilution of primary antibody. This was incubated on a nutating platform for 2h at room temperature. After incubation, beads were washed twice in Digitonin Buffer (20 mM HEPES pH 7.5, 150 mM NaCl, 0.5 mM Spermidine, 1x Roche Complete Protease Inhibitor no EDTA, 0.05% Digitonin and 0.1% bovine serum albumin), then incubated with pA-MN (600 μg/ml, 1:200, either homemade or a gift from Steven Henikoff) in Digitonin Buffer for 1 hour at 4°C. After incubation, beads were washed twice, resuspended in 150μl of Digitonin Buffer, and equilibrated to 0°C before adding CaCl_2_ (2mM) and incubating for 1 hour at 0°C. After incubation, 150μl of 2X Stop Buffer (200 mM NaCl, 20 mM EDTA, 4 mM EGTA, 50 μg/ml RNase A, 40 μg/ml glycogen), was added. Beads were incubated for 30 minutes at 37°C and then pelleted at 16,000g for 5 minutes at 4°C. Supernatant was transferred, mixed with 3 μL 10% SDS and 1.8U Proteinase K (NEB), and incubated for 1 hour at 50°C, shaking at 900rpm. After incubation, 300μl of 25:24:1 Phenol/Chloroform/Isoamyl Alcohol was added, solutions were vortexed, and transferred to Maxtrack phase-lock tubes (Qiagen). Tubes were centrifuged at 16,000g for 3 minutes at room temperature. 300μl of Chloroform was added, solutions were mixed by inversion, and centrifuged at 16,000g for 3 minutes at room temperature. Aqueous layers were transferred to new tubes and DNA isolated through Ethanol precipitation and resuspended in 10mM Tris-HCl pH 8.0 (ThermoFisher). CUT&RUN libraries were prepared following the SMARTer ThruPlex TAKARA Library Prep kit with small modifications. For each sample, double stranded DNA (10μl), Template Preparation D Buffer (2μl), and Template Preparation D Enzyme (1μl) were combined and End Repair and A-tailing was performed in a Thermocycler with a heated lid (22°C, 25 min; 55°C, 20 min). Library Synthesis D Buffer (1μl) and Library Synthesis D Enzyme (1μl) were subsequently added, and library synthesis was performed (22°C, 40min). Immediately after, Library Amplification D Buffer (25μl), Library Amplification D Enzyme (1μl), Nuclease-free water (4μl), and a unique Illumina-compatible indexed primer (5μl) were added. Library amplification was performed using the following cycling conditions: 72°C for 3 min; 85°C for 2 min; 98°C for 2 min (denaturation); 4 cycles of 98°C for 20 s, 67°C for 20 s, 72°C for 10 s (addition of indexes); 14 cycles of 98°C for 20 s, 72°C for 10 s (library amplification). Post-PCR cleanup was performed on amplified libraries with a SPRIselect bead 0.6X left/1x right double size selection then washed twice gently in 80% ethanol and eluted in 10-12μl 10mM Tris pH 8.0. 1:50 Dilutions of primary antibodies against the following active and repressive histone modifications were used: H3K4me3 (Active Motif, 39159), H3K27ac (Abcam, ab4729), H3K9me3 (Abcam, ab8898), H3K27me3 (Cell Signaling, 9733T). Primary antibodies against the following TFs were used: GATA6 (R&D Systems, AF1700), NANOG (Active Motif, 61419), SOX2 (Millipore, ab5603), OCT4 (abcam, ab19857), GATA4 (Santa Cruz Biotech, sc25310), SOX17 (R&D Systems, AF1924).

### CUT&RUN in blastocysts

Blastocysts were collected from 4-5 weeks old C57BL/6N females six days after PMSG/HCG injections. Blastocysts were used right away to profile E3.5 embryos, after removing the zona pellucida using acid tyrode’s solution. CUT&RUN was performed as described above, with significant modifications, using adaptations described by Saiz et al (Saiz et al., 2016a) for immunostainings. Embryos were manipulated in 1% agar-coated 4-well dishes (ThermoFisher scientific, Nunc(tm) 4-Well Dishes for IVF) throughout the protocol. Briefly, blastocysts were fixed using 400 μl of 0.1% formaldehyde prepared in PBS for 10 mins at room temperature, protected from light. After 10 mins, 40 μl of 1.42M glycine was added to quench PFA and embryos were incubated at room temperature for 5 mins after gentle mixing, following which they were washed once in PBS and once in digitonin buffer, by sequentially passing them through wells containing the different buffers. Embryos were then incubated in 300 μl of antibody buffer (GATA6 at 1:50) on a nutator at 4°C for 1.5h, following which they were washed thrice in digitonin buffer and then incubated in 300 μl of pA/G-MNAse solution. After incubation in pA/G-MNase, embryos were again washed thrice and then collected in a 1.5 ml microfuge tube containing 150 μl of digitonin buffer. After incubating at 0°C for 10mins, MNase was activated as described in the CUT&RUN section. The rest of the protocol was followed as described for cells using 20 amplification cycles. For the first replicate, 120 E3.5s blastocysts were used, and 160 E3.5s in the second replicate (depicted in **Fig 4F, 4G)**

### CUT&TAG

Cut&Tag was performed as previously described (Kaya-Okur et al., 2020; Kaya-Okur et al., 2019) with small modifications. mES cells were dissociated using Accutase (Sigma), counted, and 100,000 cells/replicate were pelleted at 600g for 3 minutes at room temperature. Cells were resuspended in Wash Buffer (20 mM HEPES pH 7.5, 150 mM NaCl, 0.5 mM Spermidine, 1x Protease inhibitor cocktail), and pelleted at 600g for 3 minutes at room temperature. BioMag® Plus Concanavalin A beads (Bangs Laboratories) were equilibrated in Binding Buffer (20mM HEPES pH 7.5, 10mM KCl, 1mM CaCl_2_, 1mM MnCl_2_). Cells were resuspended in Wash Buffer, mixed with a slurry of equilibrated Concavalin A coated magnetic beads, and rotated for 10 minutes at room temperature. Per 100,000 cells, a 10μl bead slurry was used. Beads were resuspended in Wash Buffer containing 2mM EDTA, 0.1% bovine serum albumin, 0.05% Digitonin, and 1:50 dilution of primary antibody and incubated on nutating platform for 2h at room temperature. After incubation, beads were washed twice in Digitonin Buffer (20 mM HEPES pH 7.5, 150 mM NaCl, 0.5 mM Spermidine, 1x Protease inhibitor cocktail, 0.05% Digitonin), then incubated with pA-Tn5 (6.7uM, 1:100) in Dig-300 Buffer (20 mM HEPES pH 7.5, 450 mM NaCl, 0.5 mM Spermidine, 1x Protease inhibitor cocktail, 0.01% Digitonin) for 1 hour at room temperature. After incubation, beads were washed twice, resuspended in 300pl of Dig-300 Buffer containing 1mM MgCl_2_, and incubated for 1 hour at 37°C.

After incubation, 10μl 0.5 EDTA (Sigma), 3μl 10% SDS, and 2.5μl 20mg/ml proteinase K (NEB) was added. Solutions were vortexed and incubated for 1 hour at 50°C, shaking at 900 rpm. After incubation, DNA separation and purification were performed as in CUT&RUN. To amplify libraries, DNA (20μl) was mixed with 5X Q5 reaction buffer (10μl), Q5 polymerase (0.5μl), dNTPs (1 μl), nuclease free water (16μl), and an equimolar mixture of an universal i5 and a uniquely barcoded i7 primer (2.5μl), using different barcodes for each sample. The sample was placed in a Thermocycler with a heated lid using the following cycling conditions: 72 °C for 5 min (gap filling); 98 °C for 30 s; 14 cycles of 98 °C for 10 s and 63 °C for 30 s; final extension at 72 °C for 1 min and hold at 8 °C. Post-PCR clean-up was performed with a SPRIselect bead 0.6X left/1x right double size selection then washed twice gently in 80% ethanol and eluted in 10-12 μl 10 mM Tris pH 8.0. Multiplexed libraries were pooled and paired-end sequenced (2 × 50 bp) on an Illumina HiSeq2500. 1:50 dilutions of primary antibodies against H3K4me3 (Active Motif, 39159), H3K27ac (Abcam, ab4729), H3K9me3 (Abcam, ab8898), H3K27me3 (Cell Signaling, 9733T) were used.

### Hi-C and Capture-C

Hi-C (4dnucleome.org protocols) and Capture-C (Downes et al., 2021) libraries were prepared following previously published protocols with minor modifications. Briefly, 1 million cells per sample were trypsinized, washed in growth media and fixed for 10 minutes at room temperature while rotating with 1% formaldehyde (Thermo: 28908) in 1ml of HBSS media. To stop fixation, Glycine was added at final concentration of 0.13M and incubated for 5 minutes at RT and 15 minutes on ice. Cells were then washed once in cold PBS, centrifuged at 2500g 4°C for 5 mins (these centrifugation conditions were used for all washes following fixation) and pellets frozen at -80°C. Thawed cell pellets were incubated in 1ml lysis buffer (10mM Tris-HCL pH8, 10mM NaCl, 0.2% Igepal CA-630, Roche Complete EDTA-free Sigma #11836170001). Following lysis, cells were dounced for a total of 40 strokes with the “tight pestle” and then washed in cold PBS. For DpnII digest, cells were resuspended in 50μl 0.5% SDS and incubated at 62°C for 10 minutes. Then 150μl of 1.5% Triton-X was added and cells incubated for 15 minutes at 37°C while shaking at 900rpm. 25μl of 10X DpnII restriction buffer (NEB) was added, and cells further incubated for 15 minutes while shaling. 200U of DpnII (NEB R0543M) were then added and incubated for 2hours, then 200U more and incubated overnight. Next morning 200U more were added and incubated for 3h (total 600U of DpnII). DpnII was inactivated at 62°C for 20 minutes. Biotin fill-in was done by incubating cells with a mixture of 4.5 μl dCTP dTTP and dGTP at 3.3 mM, 8μl klenow polymerase (NEB M0210L) and 37.5μl Biotin-14-dATP (Thermo 19524016) for 4h at RT while shaking at 900rpm for 10 seconds every 5 minutes. Ligation was done overnight at 16°C also rotating at 900rpm for 10 seconds every 5 minutes by adding 120μl of 10X ligation buffer (NEB), 664μl water, 100μl 10% Triton-X, 6μl BSA 20mg/ml, and 2μl T4 ligase (NEB cat #M0202M). For Capture-C, biotin fill-in step was skipped and 50μl more of water was added to the ligation mix. Crosslink removal was done overnight with 50μl of proteinase K in 300μl of following buffer (10mM Tris-HCl pH8.0, 0.5M NaCl, 1%SDS) while shaking at 1400rpm at 65°C. Following Sodium Acetate and 100% Ethanol -80°C precipitation, DNA was resuspended in 50μl 10mM Tris HlL for Hi-C or 130μl for Capture-C. Sonication for Hi-C was done using Covaris onetube-10 AFA strips using the following parameters for a 300bp fragment size (Duration: 10secs, repeat for 12 times, total time 120 secs, peak power-20W, duty factor 40%, CPB-50). Sonication for Capture-C was done using Covaris AFA microtubes 130 with following settings for a fragment size of 200bp fragments (Duration: 225Secs, peak power-75W, duty factor 25%, Cycles per Burst-1000). Sonications were performed in a Covaris ME220 sonicator. Sonicated material was then size selected using SPRI beads with the following ratios: 0.55X and 1X for Capture-C and 0.55X and 0.75X for Hi-C. Hi-C material was then bound, washed and recovered to 150μl Streptavidin C1 beads (Thermo 65002) per sample following manufacturers recommendations. Bead-bound DNA was ressuspended in 50μl 10mM Tris HCl. Library preparation was done using Kapa Hyper Prep KK8502 kit. 10μl of End-repair buffer and enzyme mix were added to resuspended beads and incubated for 30 minutes at RT and then 30 minutes at 65°C. 1μl of 15mM annealed-Illumina adaptors, containing an universal p5 and an indexed p7 oligo, were then incubated with a mixture containing 40pl of ligase and ligation buffer at RT for 60 minutes. Libraries were then amplified using 4 reactions per sample for a total of 200μl and 10 cycles, as recommended by manufacturer. For Capture-C, following sonication and size selection, 1μg of template material was resuspended in 50μl of 10mM Tris and used for library prep with 10μl of End-Repair reaction. 5μl of 15mM annealed -Illumina adaptors were ligated to the Capture-C material. Using a total volume of 100μl, library was amplified by PCR using 6 cycles. For capture, 1μg of Capture-C library per sample was mixed with mouse COT1 DNA and universal as well as index-specific blocking oligos from SeqCap EZ HE-oligo (Roche). 4.5μl pool of biotinylated probes (xGen Lockdown Probe Pools from IDT), with each probe at 0.4fmol/μl targeting the promoters of our loci of interested were added to this mixture and incubated for 3 days at 47°C. Following binding to Streptavidin C1 beads, material was washed as recommended by the SeqCap EZ Hybridization and Wash Kits. Following washes material was amplified by PCR using Kapa polymerase and 14 cycles. Material from different samples was then combined and 1μg of pooled libraries was recaptured in a single reaction and amplified with 8 cycles.

### ChlP-seq

ChlP-seq was performed using the ENCODE protocol with few modifications. Cells were harvested by trypsinization and counted. Approximately 30 million cells were fixed by resuspending them in 1% formaldehyde in HBSS (1ml/million cells). Cells were incubated for 10 mins at room temperature, protected from light, on a nutator. After 10 mins, 1.42M glycine (100μl/ml of fixative) was added and cells were mixed by inversion and incubated for 5 mins at room temperature to quench PFA. Cells were then pelleted at 2000g, at 4°C for 5 mins and washed twice in 1X TBS (50 mM Tris-Cl, pH 7.5, 150 mM NaCl). Cell pellets were then flash frozen and stored at -80°C. To perform ChIP, pellets were thawed on ice for 10 mins and lysed in Farnham lysis buffer (5mM PIPES PH 8.0, 85mM KCl, 0.5% NP-40, supplemented with protease inhibitors). To lyse cells, pellets were reconstituted in 1ml buffer/ 10 million cells and incubated on ice for 10 mins, following which they were dounced and then centrifuged at 2000g, at 4°C for 5 mins. The pelleted nuclei were then resuspended in 500μl RIPA (50 mM Tris-HCl pH 8.0, 150 mM NaCl, 2 mM EDTA pH8, 1% NP-40, 0.5% Sodium Deoxycholate, 0.1% SDS, supplemented with ROCHE Complete protease inhibitor tablets (no EDTA), mixed gently by pipetting, and incubated on ice for 10 mins to achieve complete lysis. Extracted chromatin was then sonicated using Bioruptor (10 cycles of 30’ on and 30’, paused on ice for 5mins, repeated 10 cycles of 30’ on and 30’). Sonicated material was then spun down at 20000g, at 4°C for 15 mins. Supernatant was collected in fresh tubes and chromatin was quantified using Qubit high sensitivity DNA kit. 10μg of chromatin was used for each ChIP reactions, in a total reaction volume of 1ml, adjusted with RIPA. For each sample, 30 μl of Protein A/G beads preconjugated to 2 μg antibody (by incubation of antibody with washed beads resuspended in 500μl RIPA for 3h) was added to the reaction and chromatin was capture by overnight incubation at 4°C, on a rotating platform. The next day, captured chromatin was washed successively as follows: once in low-salt wash buffer (0.1% SDS, 1% Triton X-100, 2 mM EDTA, 20 mM Tris-HCl pH 8.0, 150 mM NaCl), twice in high-salt wash buffer (0.1% SDS, 1% Triton X-100, 2 mM EDTA, 20 mM Tris-HCl pH 8.0, 500 mM NaCl), twice in lithium chloride wash buffer (250mM LiCl, 1% NP-40, 1% Sodium Deoxycholate, 1 mM EDTA, 10 mM Tris-HCl pH 8.0), and twice in 1X TE (10 mM Tris-HCl pH 8.0, 1 mM EDTA). Bound chromatin was then eluted in 200 μl freshly prepared direct elution buffer (10mM Tris-HCl pH8, 0.3M NaCl, 5mM EDTA pH8, 0.5% SDS) supplemented with 2 μl RNAse A (10 mg/ml), by incubating on a thermomixer at 65°C, at 800 rpm, overnight. The next day, the beads were discarded, and the supernatant was incubated with 3 μl of Proteinase K (20mg/ ml) at 55°C, 1200rpm, for 2h to reverse crosslinks. DNA was then eluted in 20 μl deionized water using the Zymo DNA Clean and Concentrator kit (Zymo, D4033). 10 μl of eluted DNA was used for library preparation exactly as described for CUT&RUN. 12 amplification cycles were used per sample, and library was sequenced on HiSeq2500 using PE50.

### RNA-seq data analysis

RNA-seq analysis including principal component analysis and identification of differentially expressed genes was performed using LCDB workflow (github.com/lcdb/lcdb-wf). Briefly, the data was first assessed for quality control using fastQC v0.11.8 (bioinformatics.babraham.ac.uk/projects/ fastqc/) to look for sequencing quality issues and trimmed for quality using cutadapt v2.7 (Martin, 2011). No significant quality issues were detected. The RNA-Seq data was then mapped to the mouse genome version GRCm38.p6 using HISAT2 v2.1.0 (Kim et al., 2015). Stats for number of reads and peaks can be found in **Table S1**. Expression levels were estimated at gene-level with featureCounts (subread v1.6.4) (Liao et al., 2014) using the GENCODE version m18 annotation. Differential expression analysis was performed using DESeq2 v1.22.1 (Love et al., 2014) using the ‘normal’ log-fold-change shrinkage. Statistical significance was defined as false discovery rate (FDR) < 0.1 and |log2FoldChange| >= 2 (4-fold difference). The count data was also normalized using the variance stabilizing transformation for use in visualization purposes. Genes with a log_2_fold change higher than 2 or lower than -2, and adjusted p value lower than 0.1 were identified as differentially expressed. Each time point was compared to previous time point and to 0h to identify all genes differentially expressed during differentiation. For plotting transcript levels at different datapoints for selected genes, DE-Seq2 normalized values were used. De-Seq2 normalized values were also used to plot changes in gene expression during differentiation in heatmaps using row zscores for each gene calculated in R. We also calculated the level of expression of the transgene compared to the endogenous *Gata6* locus. The *Gata6* transgene encodes only the coding sequence of *Gata6*. Thus, reads coming from the coding region of the gene originate both from endogenous *Gata6* and the transgene, while untranslated regions (e.g. 3’UTR) are expressed only by endogenous *Gata6*. We utilized this information to calculate the expression levels of the transgene. First, we calculated the number of reads mapping to the 3’UTR and the CDS in the last exon of *Gata6*. The counts were normalized to library size (per million) and sequence length (per kb) to avoid sequencing depth and feature length biases. The transgene expression, *T*, was then calculated as the difference between the normalized CDS and UTR expression, while endogenous expression, *E*, was taken to be the mean of the normalized 3’UTR expression at 96h (n = 4). Finally, the relative expression of the transgene was calculated as *T/E*.

### Single-cell RNA-seq analysis

Publicly available single-cell RNA-Seq data from (Nowotschin et al., 2019) was downloaded as a cell × count matrix. The cell clustering determined in the study was downloaded and the count matrix was subset to only the relevant timepoints (embryonic day E3.5 and E4.5). For use in our study we renamed and combined some clusters based on cell identities as EPI, PrE or ICM: E3.5:1 = E3.5_ICM, E3.5:0 + E3.5:4 = E3.5_PrE, E3.5:2 = E3.5_EPI-E4.5:0 + E4.5:1 = E4.5_PrE, E4.5:2 = E4.5_EPI, E4.5:3 = E4.5_TE. Next, we reanalyzed the data using the Seurat R package v3.1.2 using the standard workflow to find conserved markers and differentially expressed markers. For differential expressed markers we specifically compared PrE and EPI cells at each time-point (E3.5 and E4.5) and also E4.5_PrE and E3.5_EPI. The list of differentially expressed genes between PrE and Epi was also compared to the list reported by Mohammed and colleagues (Mohammed et al., 2017) and only genes present in both datasets was kept for further analysis. To perform a comparison between bulk and single-cell RNA-Seq data, we performed the following procedure. First, we extracted single-cell counts from cells belonging to E3.5 and E4.5 stages based on cell metadata. Next, we normalize the data to library size (per million) at cell level and performed log transformation after adding pseudocount of 1 (log1p). This is similar to the logNormalize() procedure implemented in the Seurat R package. We then averaged cells within each representative cluster. This yielded mean normalized data corresponding to each cell stage: E3.5 - ICM, PrE, EPI; E4.5 - PrE, EPI, TE. For bulk RNA-Seq, we performed variance stabilizing transformation from the DESeq2 R package to normalize count data and calculated a mean across biological replicates for each time-point. Next, for comparing such disparate sets of data, the technical variability between bulk and single-cell data needed to be regressed out. To this end, we then fit a model: *expression ∼ group*, where the factor *group* was either *single-cell* or *bulk*. The residuals from this modeling procedure was then used to generate a PCA plot.

### ATAC-seq analysis

The ENCODE ATAC-seq pipeline (github.com/ENCODE-DCC/atac-seq-pipeline) was used to process ATAC-seq and to obtain bigwig files for visualization in IGV and to produce heatmaps or signal profiles using Deeptools. For this, the bigwig files generated using signal p value and with the 2 replicates pooled were chosen. For downstream analysis, we used the IDR-identified peaks for each condition calculated from the 2 available replicates at each time point as these were the most stringent peaks identified. Diffbind (Ross-Innes et al., 2012) was used to plot PCA and identify peaks with differential accessibility during differentiation by comparing 48h to 0h. Stats for number of reads and peaks can be found in **Table S1**. For comparisons, the DESEQ2 option was used and an adjusted pvalue of 0.0001 and log2 fold change of 2(or -2) was used as threshold of significance. All timepoints were compared against 0h. Following classification of ATAC peaks into stable, increased, or decreased throughout 48h of differentiation we used GREAT (McLean et al., 2010) to further identify which of these peaks were Epi or PrE specific. For this, peaks were assigned to nearest gene with a maximum distance of up to 50kb. A combination of Deeptools (Ramirez et al., 2016) and bedtools (Quinlan and Hall, 2010) was used for plotting heatmaps and summary profiles. Deeptools profiles and heatmaps were plotted using peak center as the single reference point, profiles were plotted using mean signal. ATAC-seq nucleosomal signal over TF motifs was plotted using the ATACseqQC (Ou et al., 2018) pipeline by merging both replicates for the same time point. Only nucleosomal signal (fragments larger than 180bp) is shown. For this, we used FIMO (Bailey et al., 2009) to identify TF motifs at peaks of interest identified by CUT&RUN. FIMO was run using a pvalue threshold of 0.001 and with the following JASPAR motifs: GATA6 (PB0023.1), NANOG (UN0383.1). To plot ATAC footprinting signal over HNF1B motifs (MA0153.1) we used the TOBIAS pipeline (Bentsen et al., 2020) with merged ATAC-seq replicates. ATAC signal was plotted over all motifs identified as bound at 48h by this pipeline.

### CUT&RUN and CUT&TAG analysis

Paired-end 50bp reads were processed using Bowtie2 (Langmead and Salzberg, 2012), with the following options (-N 1 --local --very-sensitive-local --no-unal --no-mixed --nodiscordant --phred33 -I 10 -X 700 -x). Reads that mapped to ENCODE mm10 blacklist regions were removed using Samtools (Li et al., 2009). Piccard (broadinstitute.github. io/picard/) was then used to identify non-duplicated reads. Duplicate reads were removed for libraries generated from blastocysts but kept for all others. Diffbind was used to analyze replicates by plotting pearson correlation of signal found over identified peaks (**Fig S7**). For IGV visualization and for plotting heatmaps and profiles using Deeptools, replicate signal was merged and normalized. Epi TFs and CUT&TAG were normalized using total reads. Stats for number of reads and peaks can be found in **Table S1**. For Epi TFs, signal normalization was used using reads that mapped to *E*.*coli* and *S*.*cereviseae* genomes. For TF signal, only fragments smaller than 120bp were used while for histone modifications, fragments larger than 150bp were selected. MACS was used to identify regions of enrichment using the narrow option for TFs and the broad option for histone modifications. Peak calling was done using as a control CUT&RUN samples generated using rabbit antirabbit antibody. Peaks overlapping with ENCODE blacklist regions were removed from further downstream analysis. A combination of deeptools and bedtools was used for plotting heatmaps and summary profiles. Characterization of GATA6 peak types was done considering the top 10000 peaks at each time point according to adjusted q value calculated by MACS. This valued was chosen based on previous comparison to published GATA6 ChIP-seq data (Wamaitha et al., 2015). Peaks were considered as Early if present in the top 10000 peaks at any timpoint between 2 and 8h for a total of 14906 early peaks. If present only at 48h peaks were identified as Late (total 5850). Early peaks that intersected with ATAC peaks at 0hours were considered as Early Open at 0h and the remaining Early peaks were considered Early Closed at 0h. These peaks were then associated with PrE or Epi genes using the same approach as for ATAC-seq. Peaks were further divided into proximal or distal depending on distance to TSS (less than 5kb away were considered proximal). NANOG peaks were identified similarly to GATA6 and the top 15000 peaks selected for characterization based on number of peaks detected by ENCODE data. To define which peaks overlapped with GATA6, all GATA6 peaks identified from 2,4 and 8h were used. NANOG peaks were classified as Epi or PrE specific as done for ATACseq. For analysis of NANOG peaks that appear specifically at 2h, all peaks were considered and separated into 3 clusters. Peaks that overlapped with NANOG peaks already identified at 0h and peaks that were only identified at 2h. These were separated into those that overlapped with ATAC-seq peaks at 0hours and those that didn’t. CUT&RUN analysis of Cutsite probability was performed using CUTRUNTools (Zhu et al., 2019) using the GATA6 (PB0023.1) motifs identified by FIMO over the mentioned peaks in figure. A threshold of 0.001 was used to identify motifs within peaks.

### TF motif analysis

To identify TF motifs for GATA6 and OCT4/SOX2 at peaks of interest we used FIMO with a pvalue threshold of 0.001 and with the following JASPAR motifs: GATA6 (PB0023.1), and OCT4/SOX2 (MA0142.1). Then, for each peak we calculated the number of motifs found in each peak and displayed this number as a boxplot. To assess motif strength, for all motifs identified with a pvalue lower than 0.001, we plotted the FIMO-calculated qvalue. For comparison of motifs in two different conditions we used Homer to identify known motifs enriched in one condition versus another as shown in figure.

### Hi-C and Capture-C analysis

Hi-C libraries were sequenced with paired-end reads of 51 nucleotides. Data was processed using the Hi-Cpro pipeline (Servant et al., 2015) to produce a list of valid interactions pairs. This list was converted into cool and mcool files for visualization with higlass (Kerpedjiev et al., 2018). Stats for number of reads and peaks can be found in **Table S1**. Eigenvalues were calculated using homer (Heinz et al., 2018) with 250kb bins and the runhicpca.pl script. For comparison between conditions, we used the gethiccorrDiff. pl script also at 250kb resolution. Bins were considered as belonging to different compartments if correlation coefficient was lower than 0.4. For heatmaps containing eigenvalues, bins were sorted from highest eigenvalue to lowest based on the 0h score. H3K27me3 and H3K27ac score for each bin was determined using Deeptools. Capture-C were libraries were also sequenced and processed the same way. The make_viewpoints Hicpro script was used to obtain individual Capture-C bigwig files for each replicate of each viewpoint with 1kb-sized bins. For visualization, averages of each replicate were used. P values were calculated using DESeq2 and comparing Capture-C signal of 2 replicates over overlapping 5kb windows across the regions shown in figure. Overlapping 5kb windows were built by sliding each window 500bps. Comparisons were done between 0h and either 16h or 48h. Adjusted p values lower than 0.01 were considered as threshold for significance. Horizontal bars represent windows considered as statistically significantly different for these comparisons.

## ACKNOWLEDGMENTS

We thank all members of the Unit on Genome Structure and Regulation for discussions and Nestor Saiz, Effie Apostolou, Karl Pfeifer, and Judith Kassis for comments on the manuscript. We thank NICHD’s molecular genetics core, specifically Steven Coon, Tianwei Li and James Iben. This work utilized the computational resources of the NIH HPC Biowulf cluster (http://hpc.nih.gov). We thank Lisa Price for cell sorting experiments. We thank the mouse core of NICHD specifically Jeanne Yimdjo, Victoria Gibbs and Alexander Grinberg. We are also in debt to Kathy Niakan for providing the inducible GATA6 mES cells and Steven Henikoff for pA-MNase and pA-Tn5 enzymes. This work was funded by NIH intramural project HD008975-02. All data can be downloaded from GEO under accession GSE181073.

## AUTHOR CONTRIBUTIONS

JT, DL, SF and PPR performed experiments. JT, AM, and PPR analyzed experiments. RD and PPR supervised the project. JT and PPR designed the project and wrote the paper with input from all other authors.

---

**Supplementary figure S1:**
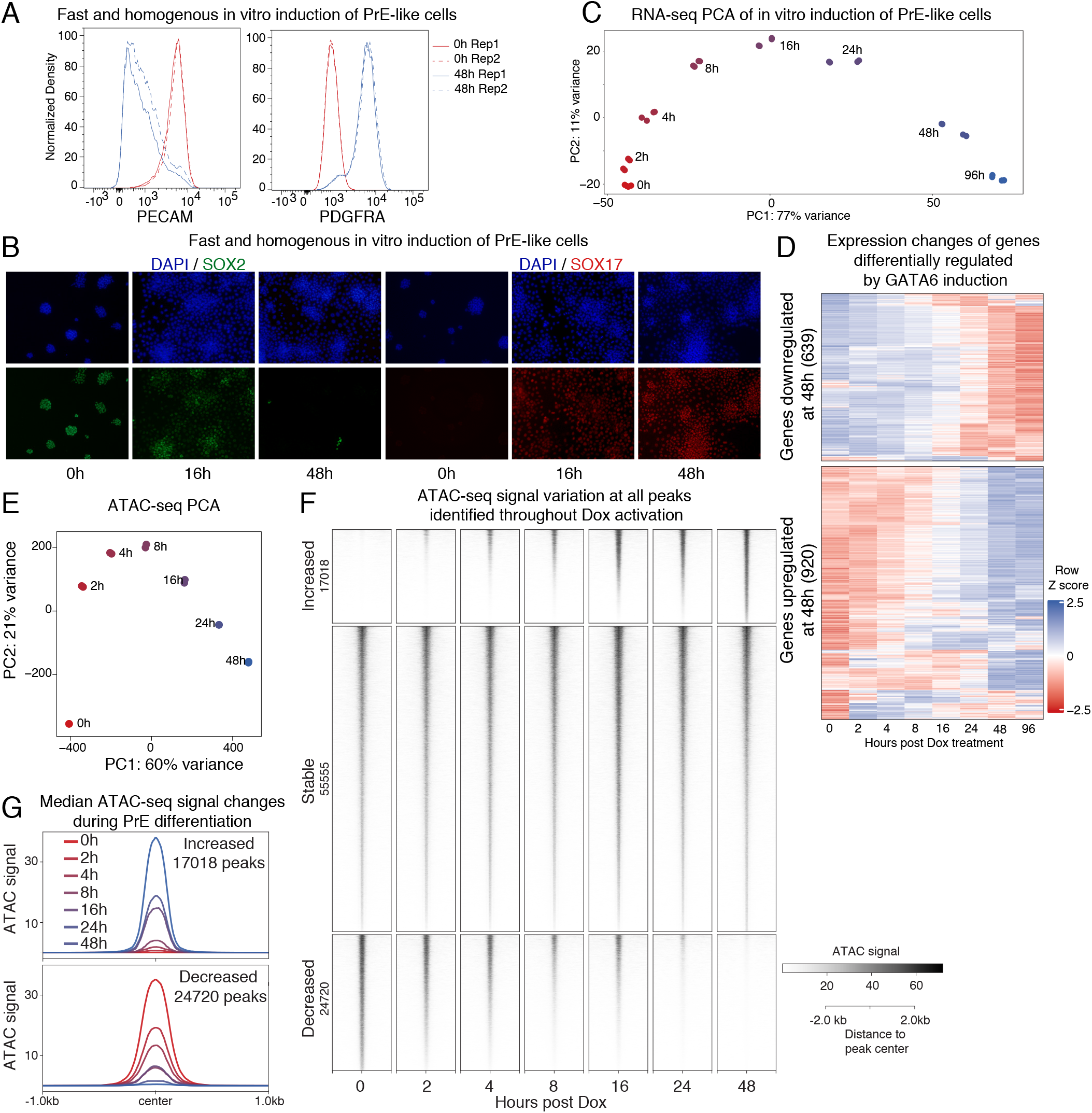
GATA6-induction drives quick transcriptional changes and remodelling of chromatin accessibility. **A** Flow cytometry analysis at 0h and 48h on cells co-stained for the pluripotency surface marker, PECAM, and the PrE-specific surface marker, PDGFRA. Differentiation was associated with homogeneous loss of PECAM and gain of PDGFRA. **B** SOX2 IF (green) showed that the Epi-state was quickly and homogeneously lost with concomitant expression gain of the PrE marker, SOX17 (red). **C** PCA plot of transcriptome changes accompanying differentiation, as profiled by bulk RNA-seq shows that transcriptional changes were quickly initiated by 2h of GATA6 expression. **D** Change in expression of all genes identified as differentially expressed during GATA6-induction represented as a heatmap of zscores for each gene. Plot in Fig 1C shows only the genes identified as differentially expressed between PrE and Epi in single cell blastocyst RNA-seq data. **E** PCA of chromatin accessibility throughout differentiation as assayed by ATAC-seq showing that just like the transcriptome, chromatin remodeling was quickly initiated by 2h of GATA6 expression. **F** Heatmap showing the changes in ATAC-seq signal at all identified ATAC-seq peaks. Both accessibility gain and loss were initiated within 2h of GATA6-induced PrE differentiation. Changes in accessibility are categorized into chromatin regions that gained accessibility (increased), lost accessibility (decreased) and regions where accessibility remained unchanged (stable, 55555 peaks) when comparing 0h to 48h. **G** Median ATAC-signal at each time point plotted at the 17018 peaks across the genome that gained accessibility during differentiation (top panel) and at the 24720 peaks that lost accessibility.

**Supplementary Figure 2.**
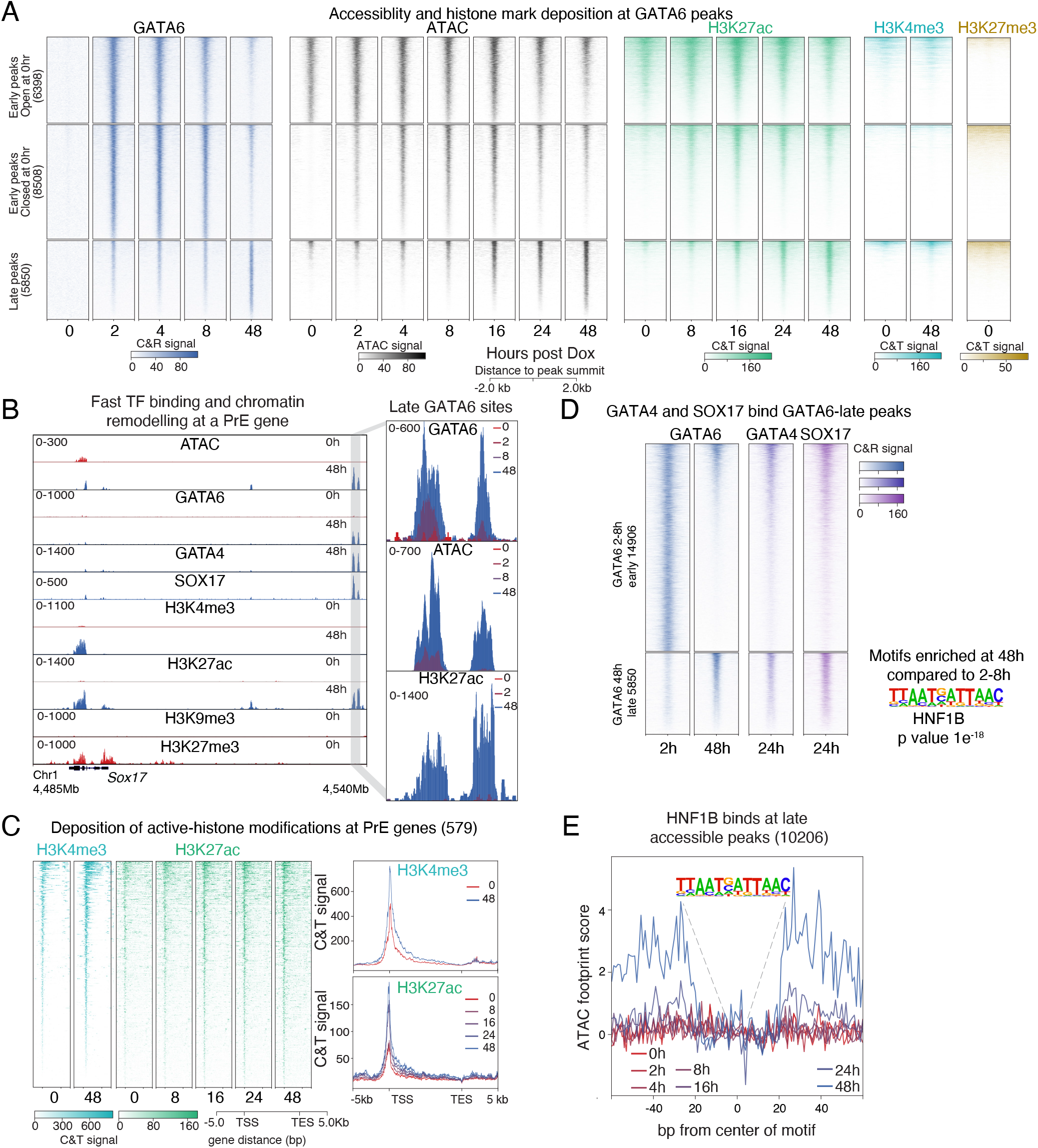
GATA6 binding leads to gains in chromatin accessibility and H3K27ac deposition. **A** Heatmap depicting changes in chromatin accessibility and distribution of histone marks at top 15000 GATA6 peaks identified across the mouse genome. GATA6 binding sites were classified into early peaks (bound by GATA6 within 8 hours of differentiation) and late peaks (bound by GATA6 only at 48 hours). Early peaks were further divided into those accessible before differentiation (Early peaks open at 0hr) and those that gained accessibility only after the induction of differentiation (early peaks closed at 0hr). **B** Browser view at the *Sox17* locus showing the change in chromatin landscape at 48 hours of differentiation. Zoomed-in view of the grey highlighted region shows that this putative *Sox17* CRE is a late GATA6 binding site that is inaccessible at 0hrs and only reached high GATA6 binding and increase in chromatin accessibility by 48 hours. **C** Heatmap showing deposition of H3K4me3 and H3H27ac at PrE gene-bodies as differentiation progresses. Median signal profile plots alongside show an increase of H3K4me3 and H3K27me3 at later time points in differentiation compared to the 0hr time-point. **D** GATA4 and SOXI7 bind most of GATA6 late-binding peaks. GATA4 and SOX17 24h CUT&RUN signal plotted at early (found between 2-8h) and late peaks (found only at 48h). GATA6 late peaks are enriched for the HNF1B motif when compared to motifs found from 2-8 hours using HOMER. **E** ATAC footprint score calculated using TOBIAS with the HNF1B motif at CREs that only become accessible at 48h. ATAC footprint score showing higher signal surrounding the HNF1B motif suggests HNF1B binding starting at 24h.

**Supplementary Figure 3.**
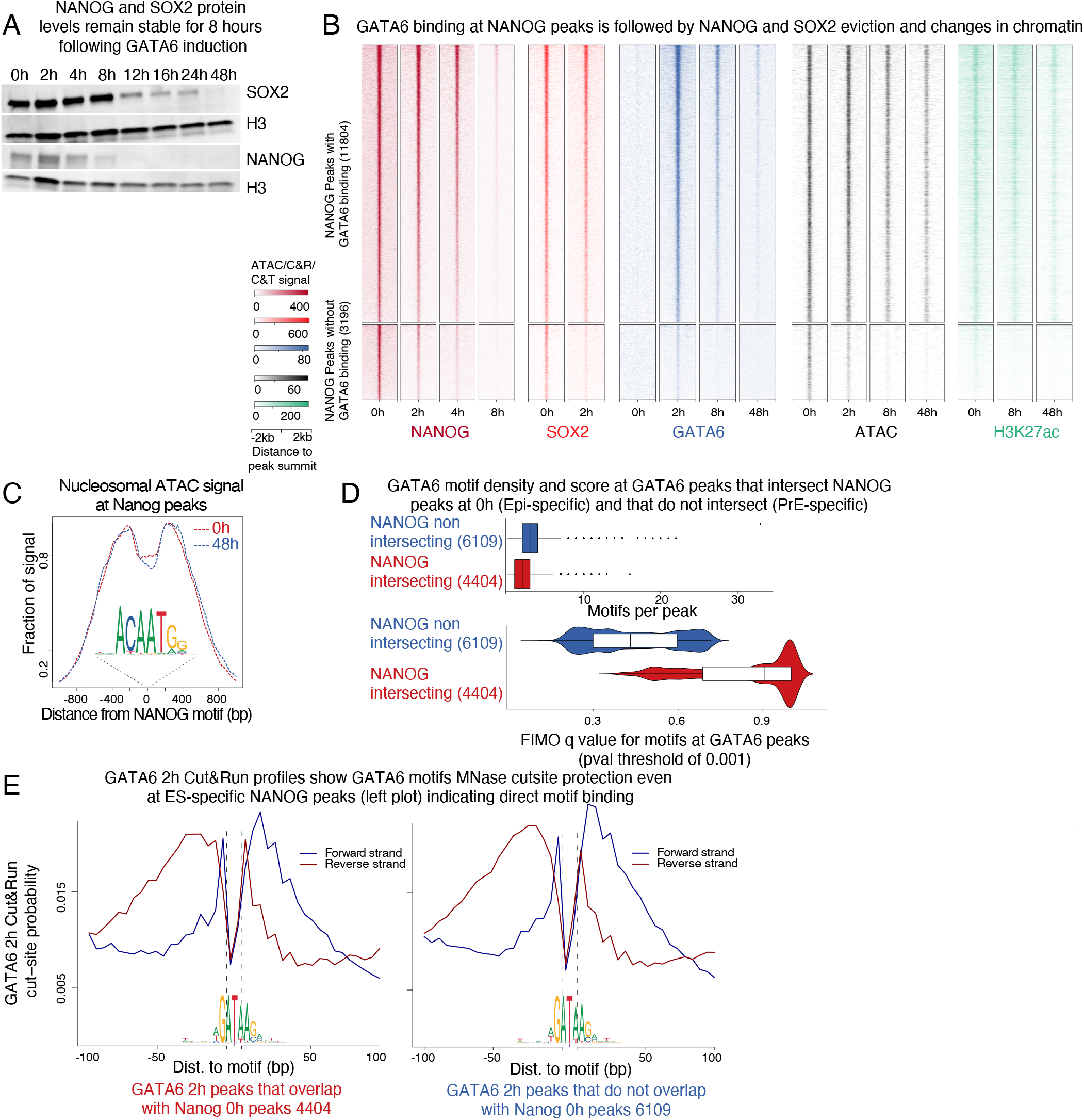
GATA6 transiently binds most NANOG-bound regions prior to inactivation. **A** Western blot of SOX2 and NANOG showing that protein levels of these two Epi-TFs is not decreased by 2h of differentiation and that NANOG is repressed faster than SOX2. **B** Heatmap depicting gradual loss of NANOG and SOX2 at NANOG peaks, concomitant with the transient binding of GATA6 at these loci, gradual loss of ATAC signal, and gradual depletion of H3K27ac with little change in H3K27me3 levels. The top 15000 NANOG peaks were divided into peaks where GATA6 overlapped with NANOG (top) and peaks where GATA6 did not overlap with NANOG. **C** Nucleosomal fraction of ATAC-seq signal as determined by ATACseqQC at the indicated timepoints over NANOG motifs at NANOG-bound peaks shows minimal reshuffling of nucleosomes even at 48 hours post differentiation implying that NANOG motif occlusion by nucleosomes did not play a determinant role in silencing of the Epi transcriptional program. **D** Density and FIMO q value score of GATA6 motifs at GATA6 peaks that NANOG did not bind at 0h (PrE-specific -blue) and peaks that were bound by NANOG at 0h (Epi-specific-red). Epi-specific peaks also contained GATA6 motifs but at lower density and weaker affinity compared to the GATA6 motifs at PrE-specific peaks. **E** Cut&RUN cutsite probability at GATA6 peaks that overlap with NANOG and are Epi CREs (left) and GATA6 motifs located in GATA6 peaks that did not overlap with NANOG and are PrE specific (right). In both cases cutsite probability was similar and symmetric displaying protection of the GATA6 motif, suggesting that GATA6 motifs are indeed occupied by GATA6 in both types of peaks.

**Supplementary Figure 4.**
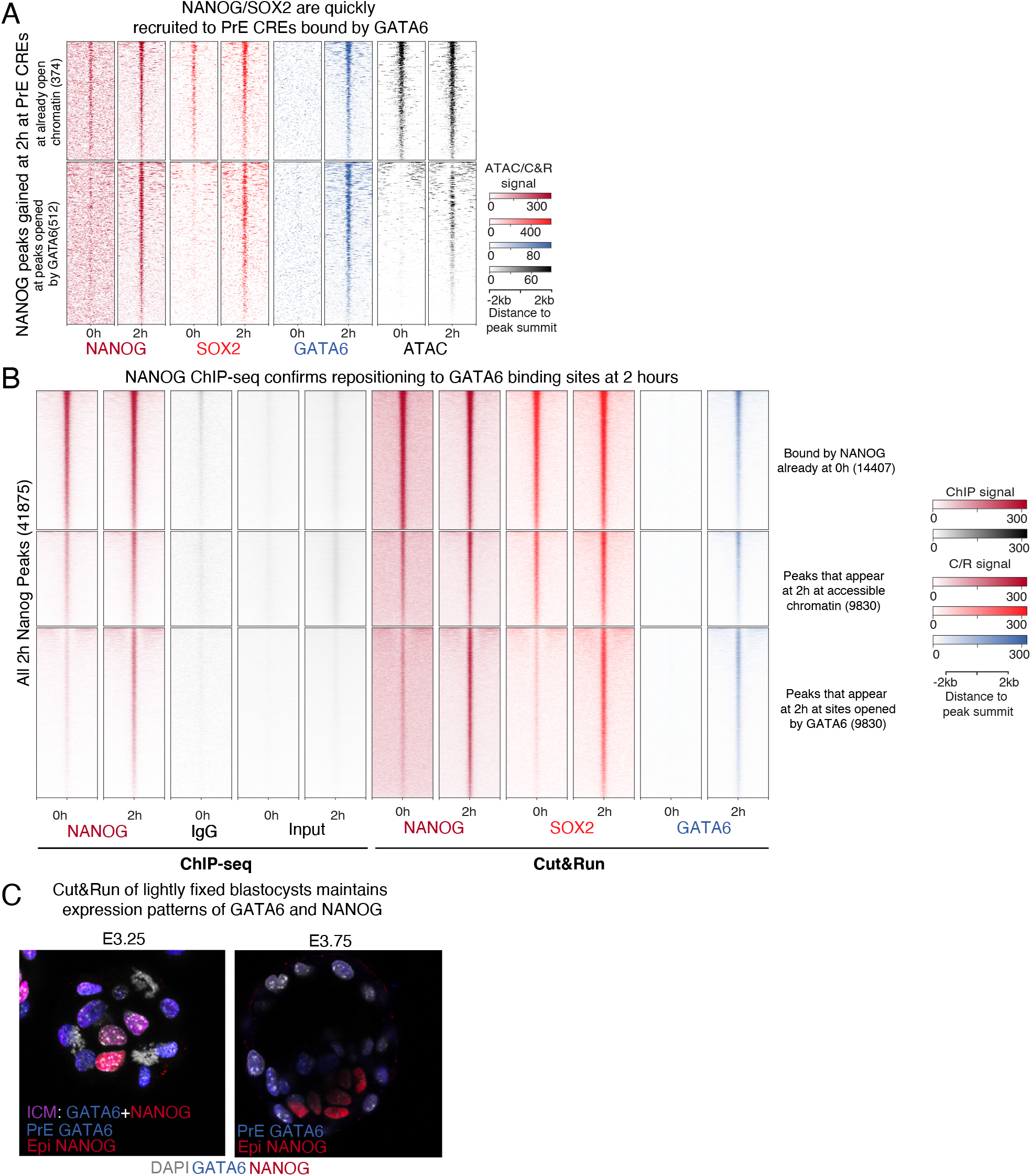
Genome wide binding of NANOG and SOX2 to GATA6 targets. **A** Heatmap showing de novo recruitment of NANOG and SOX2 at GATA6 CREs associated with PrE genes. **B** Heatmap with same clusters as in Fig.4A showing that repositioning of NANOG to GATA6 peaks seen with Cut&RUN is also observable using ChIP-seq. **C** Example of blastocysts processed with our modified CUT&RUN protocol but hybridized with fluorescently-labeled secondary antibodies. Picture shows that light fixation maintained blastocyst structure and detected changes in tissue-specific expression of both GATA6 and NANOG. At E3.25, ICM cells that expressed both TFs can be seen (purple) as well as cells already committed to either TE (white), PrE (blue) or Epi(red). At later blastocyst stages, E3.5-E3.75 only TE, Epi and PrE-commited cells are found and no ICM cells are seen.

**Supplementary Figure 5.**
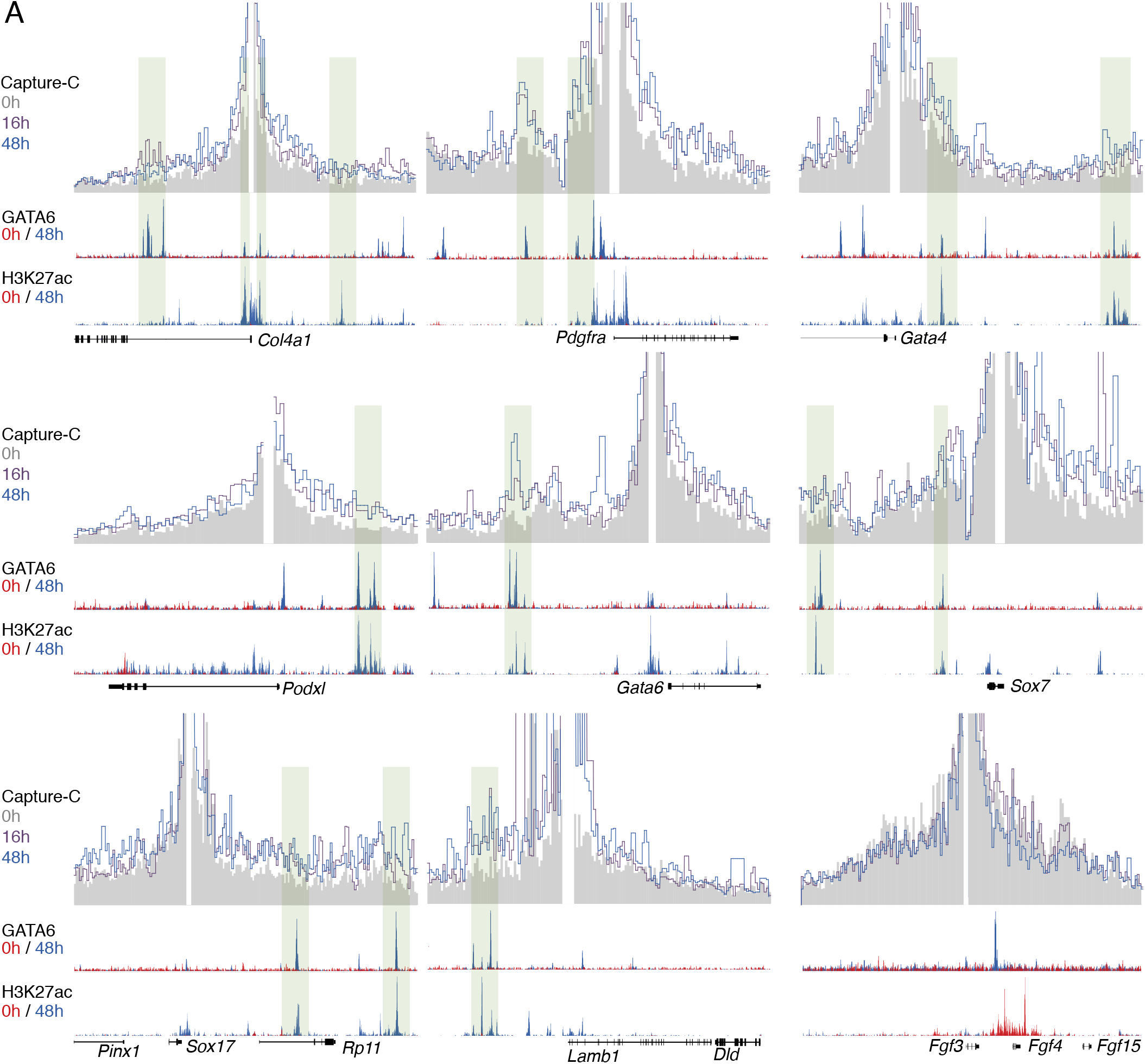
PrE genes increase interaction frequency with GATA6-induced CREs within 16 hours of differentiation. All gene promoters displayed an increase in interactions during differentiation with regions that are bound by GATA6 and that gain H3K27ac. The only exception is *Fgf3*, which is a PrE-specific gene but does not display distal CREs controlling its expression possibly explaining a lack of 3D structure reconfiguration. Capture-C data is shown as the average signal of two replicates using 1kb bins. Green shaded area represents putative CREs with increased interactions with PrE-specific genes.

**Supplementary Figure 6.**
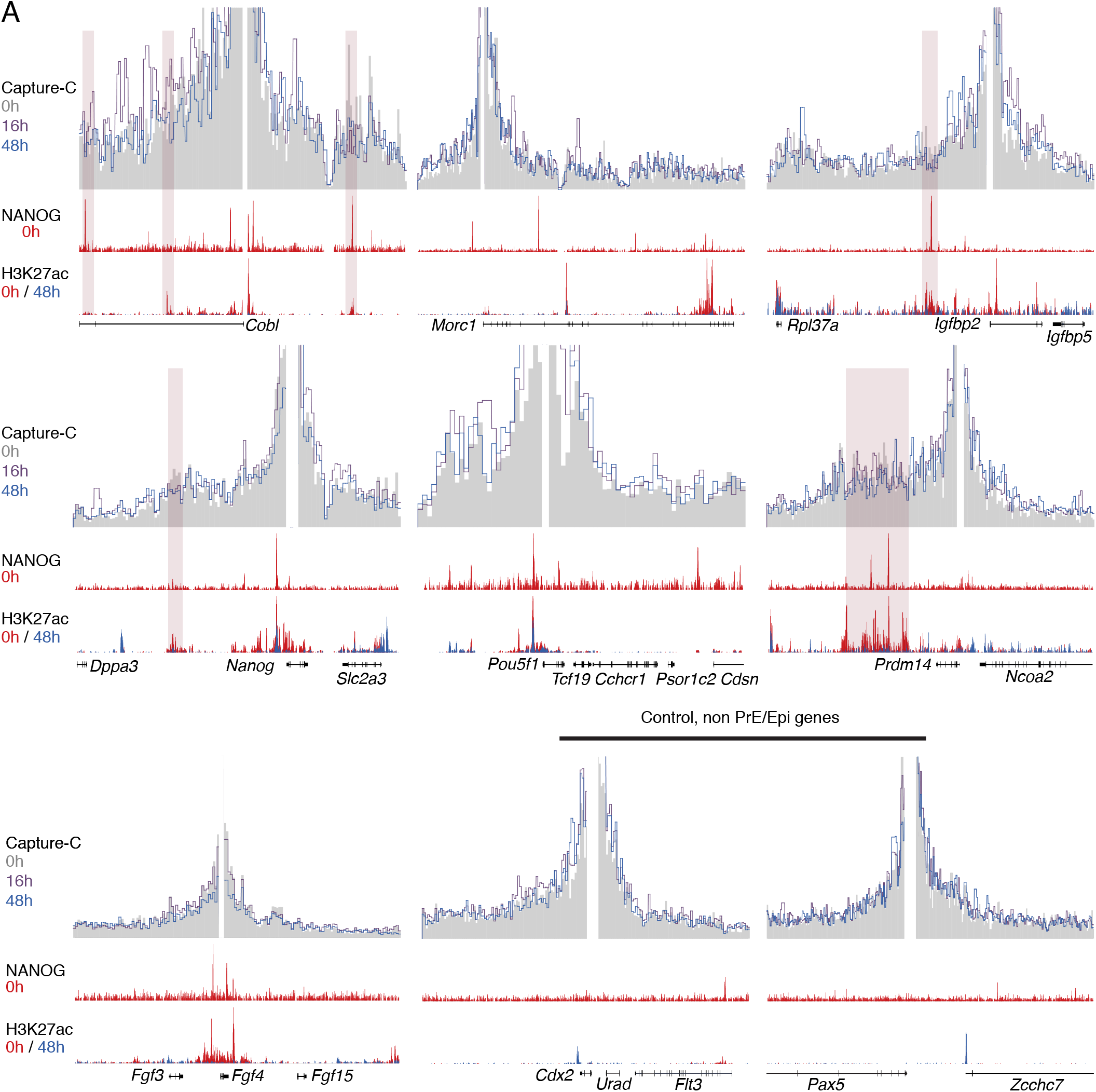
Epi genes decrease interaction frequency with Epi-specific CREs within 16 hours of differentiation. Most Epi genes show a decrease in interaction frequency during differentiation with regions that are bound by NANOG at 0h and that lose H3K27ac. The exceptions are *Morc1, Fgf4* and *Pou5f1*. None of these genes shows putative distal CREs possibly explaining higher stability of interactions. Two control genes that do not change expression during differentiation are shown and display no changes in interaction frequencies. Capture-C data is shown as the average signal of two replicates using bins of 1kb. Red shaded area represents putative CREs with decreased interaction frequency with PrE-specific genes.

**Supplementary Figure 7.**
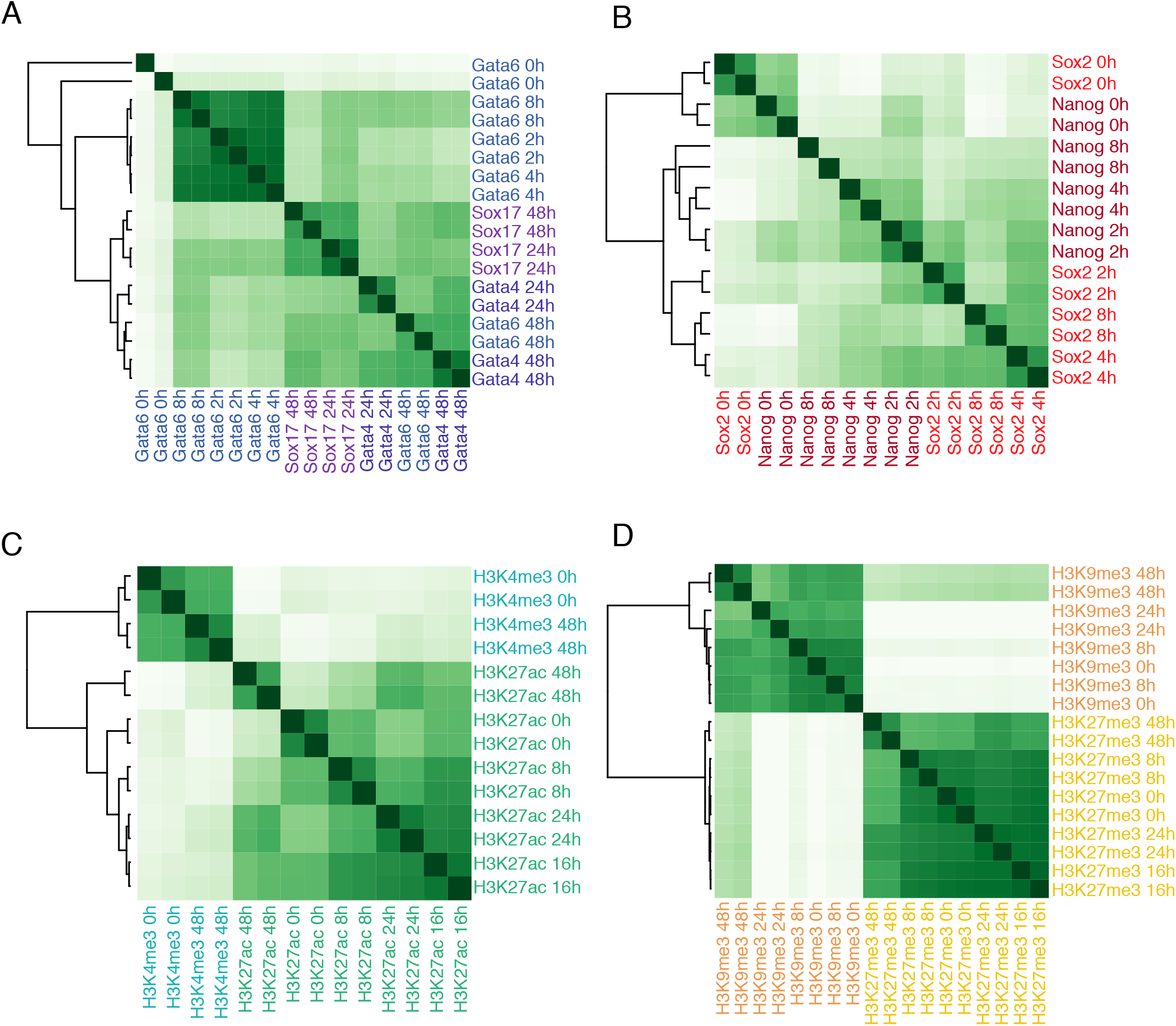
Correlation analysis between replicates of CUT&RUN and CUT&TAG datasets. The R-package DiffBind was used to compare between datasets. **A** Correlation plot between the two replicates of each PrE transcription factor profiled at the different time points. **B** Correlation plot between the two replicates of each TF profiled at the different time points. **C** Correlation plot between the two replicates of active histone marks profiled by CUT&TAG at the indicated time points and **D** Correlation plot between the two replicates of repressive histone marks profiled at the different time points.

## Notes

https://hpc.nih.gov/~GSRunit/Thompson_etal.tar.gz

### Competing Interest Statement

The authors have declared no competing interest.

